# Hierarchical Bayesian Augmented Hebbian Reweighting Model of Perceptual Learning

**DOI:** 10.1101/2024.08.08.606902

**Authors:** Zhong-Lin Lu, Shanglin Yang, Barbara Dosher

## Abstract

The Augmented Hebbian Reweighting Model (AHRM) has been effectively utilized to model the collective performance of observers in various perceptual learning studies. In this work, we have introduced a novel hierarchical Bayesian Augmented Hebbian Reweighting Model (HB-AHRM) to simultaneously model the learning curves of individual participants and the entire population within a single framework. We have compared its performance to that of a Bayesian Inference Procedure (BIP), which independently estimates the posterior distributions of model parameters for each individual subject without employing a hierarchical structure. To cope with the substantial computational demands, we developed an approach to approximate the likelihood function in the AHRM with feature engineering and linear regression, increasing the speed of the estimation procedure by 20,000 times. The HB-AHRM has enabled us to compute the joint posterior distribution of hyperparameters and parameters at the population, observer, and test levels, facilitating statistical inferences across these levels. While we have developed this methodology within the context of a single experiment, the HB-AHRM and the associated modeling techniques can be readily applied to analyze data from various perceptual learning experiments and provide predictions of human performance at both the population and individual levels. The likelihood approximation concept introduced in this study may have broader utility in fitting other stochastic models lacking analytic forms.

## INTRODUCTION

Perceptual learning is a powerful process that can significantly enhance human performance in various perceptual tasks (Dosher & Lu, 2020; Fahle & Poggio, 2002; Green, Banai, Lu, & Bavelier, 2018; Lu & Dosher, 2022; Lu, Hua, Huang, Zhou, & Dosher, 2011; Sagi, 2011; Seitz, 2017; T. Watanabe & Sasaki, 2015). It can lead to improvements in tasks such as orientation, spatial frequency, and motion direction judgements, taking performance from near chance to high proficiency (Ball & Sekuler, 1982; Fiorentini & Berardi, 1980; Poggio, Fahle, & Edelman, 1992). Contrast sensitivity can increase by over 100% (Dosher & Lu, 1998; Huang, Zhou, & Lu, 2008), and response times can decrease by approximately 40% (Petrov, Van Horn, & Ratcliff, 2011). Perceptual learning is increasingly being applied in rehabilitation and the development of perceptual expertise (Cavanaugh, 2015; L. Gu et al., 2020; Hess & Thompson, 2015; Huang et al., 2008; Huxlin et al., 2009; Levi, 2020; Lu, Lin, & Dosher, 2016; Maniglia, Visscher, & Seitz, 2021; Roberts & Carrasco, 2022; F.-F. Yan et al., 2015).

Two main theories, representation enhancement and selective reweighting, have been proposed to explain performance improvements in visual perceptual learning (Ahissar & Hochstein, 2004; Dosher & Lu, 1998, 2009b; Fahle, 1994; Karni & Sagi, 1991; Mollon & Danilova, 1996; Sotiropoulos, Seitz, & Seriès, 2011; Talluri, Hung, Seitz, & Seriès, 2015; T. Watanabe et al., 2002; T. Watanabe & Sasaki, 2015; Zhang et al., 2010). Representation enhancement suggests that perceptual learning improves performance by altering the responses or tuning characteristics of neurons in early visual cortical areas. On the other hand, selective reweighting involves the up-weighting of relevant and down-weighting of irrelevant representations from early visual cortical areas during perceptual decision without changing the representations themselves. While both processes can contribute to perceptual learning (Kourtzi, Betts, Sarkheil, & Welchman, 2005; Roelfsema, van Ooyen, & Watanabe, 2010; Seitz & Watanabe, 2005; T. Watanabe & Sasaki, 2015), selective reweighting appears to be the dominant component (Dosher & Lu, 2020). This conclusion is also supported by physiological and brain imaging studies, which indicate that early sensory representation enhancement accounts for only a small fraction of behavioral performance improvements (Ghose, Yang, & Maunsell, 2002; Schoups, Vogels, Qian, & Orban, 2001), while evidence of neural plasticity is strongest in higher visual areas (Adab & Vogels, 2011; Law & Gold, 2008; Y. Yan et al., 2014). Notably, representation enhancement remains primarily a verbal theory, and most existing computational models of visual perceptual learning are based on selective reweighting. These models aim to enhance the readout of sensory information during perceptual decision making (Dosher, Jeter, Liu, & Lu, 2013; Dosher & Lu, 1998; Jacobs, 2009; Law & Gold, 2009; Petrov, Dosher, & Lu, 2005; Poggio et al., 1992; Sotiropoulos et al., 2011; Vaina, Sundareswaran, & Harris, 1995; Weiss, Edelman, & Fahle, 1993; Zhaoping, Herzog, & Dayan, 2003).

The Augmented Hebbian Reweighting Model (AHRM; Figure 1) is the first full computational process model of perceptual learning (Petrov et al., 2005). It comprises representation, bias, feedback, and decision modules. The representation module computes activations in multiple orientation- and frequency-selective channels from stimulus images. The decision module weights and sums activations along with a bias module and yields a response on each trial. The learning module updates the weights to the decision module on every trial using augmented Hebbian learning, which moves the “late” post-synaptic activation in the decision module towards the correct response when feedback is available, and operates on the early decision activation when there is no feedback (Dosher & Lu, 2009a; Petrov et al., 2005; Petrov, Dosher, & Lu, 2006).

**Figure 1.**
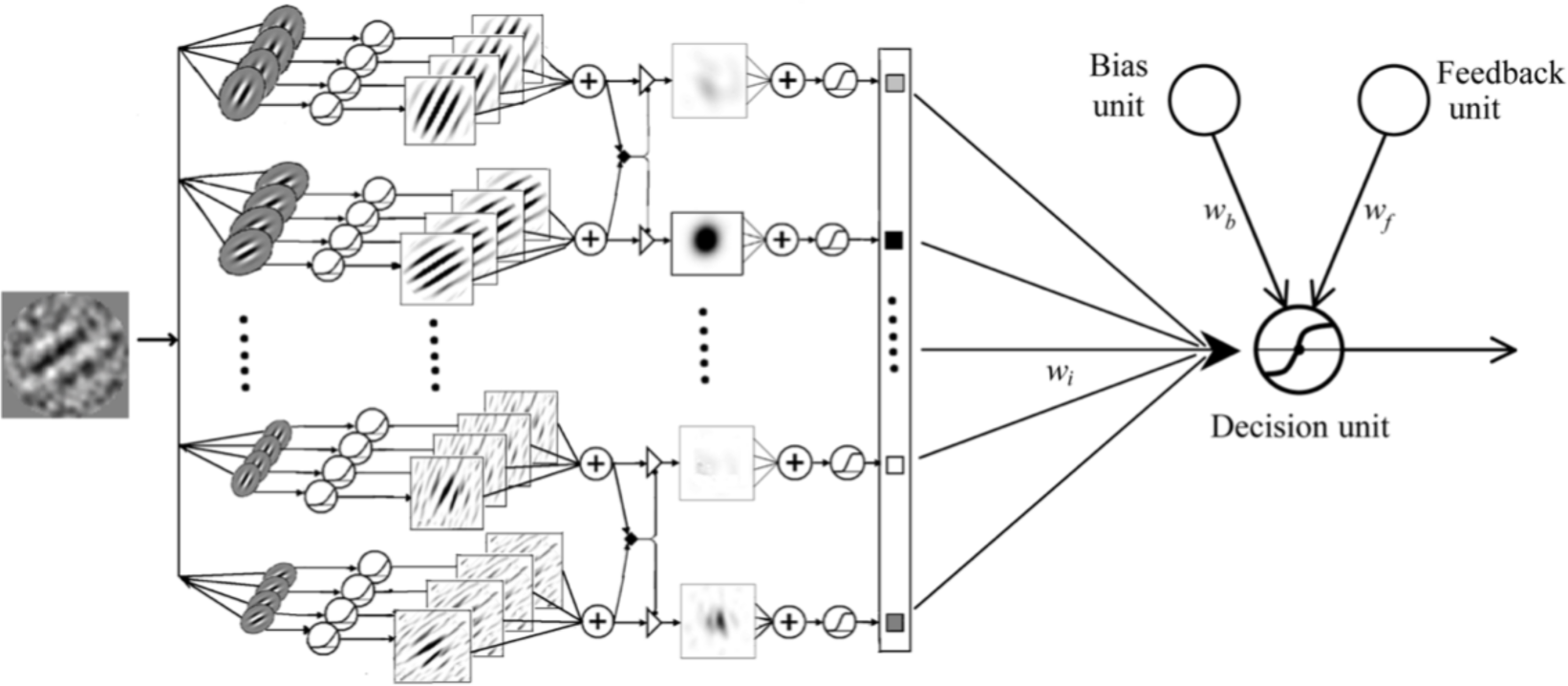
The Augmented Hebbian Reweighting Model (AHRM) with representation, bias, feedback, and decision modules.

The AHRM has successfully modeled various phenomena in perceptual learning, including perceptual learning in nonstationary environments with and without feedback (Petrov et al., 2005, 2006), basic mechanisms of perceptual learning, asymmetric transfer of learning in high and low external noise, and effects of pretraining mechanisms (Lu, Liu, & Dosher, 2010), co-learning of multiple tasks (Huang, Lu, & Dosher, 2012), interaction between training accuracy and feedback (Liu, Lu, & Dosher, 2010; Liu, Lu, & Dosher, 2012), and trial-by-trial and block feedback (Liu, Dosher, & Lu, 2014). It has also led to the development of several related models (Dosher et al., 2013; Jacobs, 2009; Law & Gold, 2009; Sotiropoulos et al., 2011; Talluri et al., 2015).

Despite its success, fitting the AHRM to data presents a significant challenge. The AHRM is a sequential stochastic learning model, that, with a given set of parameters, must be simulated to generate performance predictions with sequential trial-by-trial updates of the decision weights. Simulations typically involve running the model hundreds to thousands of times to generate average predictions and confidence bands for a given set of parameter values. For a fixed set of parameter values, each run of the model leads to a different sequence of responses and somewhat different weight changes due to stochastic trial-by-trial variations resulting from internal and external noises and different random trial sequences. Fitting the AHRM with typical curve fitting procedures (e.g., maximum likelihood, least squares, Bayesian) is not feasible because the fitting process requires simulations of many potential parameter sets (tens to hundreds of thousands). Instead, estimation of the AHRM parameters is generally done using hierarchical grid-search methods. These methods evaluate a matrix of spaced parameter values and then narrow down regions of the parameter space that are more promising, making it difficult to obtain the optimal solutions.

In this study, we introduce three modeling technologies to facilitate AHRM fitting:

1. *A Hierarchical Bayesian AHRM (HB-AHRM)*: This approach incorporates population, subject, and test levels to estimate the joint posterior hyperparameter and parameter distribution across all subjects while considering covariance within and between subjects.
2. *Vectorization with PyTensor*: Leveraging PyTensor library and GPU acceleration, these techniques drastically speed up simulations by optimizing the computation of mathematical expressions involving multi-dimensional arrays.
3. *Likelihood function approximation*: We developed an approach to approximate the likelihood function in the AHRM with feature engineering and linear regression. Based on simulated predictions of the AHRM over a large parameter grid, we encoded the functional relationship between the likelihood and parameters, greatly accelerating model computations.

Hierarchical models enable effective combination of information across subjects and groups while preserving heterogeneity (Kruschke, 2014; Rouder & Lu, 2005). These models typically consist of sub-models and probability distributions at multiple levels of the hierarchy and can compute the joint posterior distributions of the parameters and hyperparameters using Bayes’ theorem based on all available data (Kruschke, 2014; Kruschke & Liddell, 2018). Hierarchical models are valuable for reducing the variance of estimated posterior distributions by decomposing variabilities from different sources using parameters and hyperparameters (Song et al., 2020) and shrinking estimated parameters at lower levels towards the modes of higher levels when there is insufficient data at the lower level (Kruschke, 2014; Rouder et al., 2003; Rouder & Lu, 2005).

The HB-AHRM consists of three levels: population, subject, and test. In this framework, all subjects belong to a population and may, in principle, run the same experiment (called “test”) multiple times. The distributions of AHRM parameters at the test level are conditioned on the hyperparameter distributions at the subject level, which, in turn, are conditioned on the hyperparameter distribution at the population level. The HB-AHRM also includes covariance hyperparameters at the population and subject levels to capture the relationship between and within subjects.

PyTensor is a Python library used to define, optimize, rewrite, and evaluate mathematical expressions, particularly those involving multi-dimensional arrays. It combines elements of a computer algebra system and an optimizing compiler. PyTensor is particularly useful for tasks where complex mathematical expressions need repeated evaluation, and speed is critical. The library provides a loop mechanism called *scan*, which can process inputs efficiently. We used PyTensor to represent all variables in the HB-AHRM and applied the *scan* function, significantly speeding up simulations from 22.2 to 1.6 seconds for 300 repeated runs of the experiment in Petrov et al. (2005) based on one set of AHRM parameters.

Although PyTensor improved simulation speed, computing the HB-AHRM still involves evaluating of hundreds of thousands of parameter sets. Because of the tremendous computational load, we developed a method to approximate the likelihood function in the AHRM with feature engineering and linear regression. It involves simulating AHRM predictions in a large parameter grid using parallel computing with GPU processors, taking <24 hours for a 64,000 mesh grid. We then employed feature engineering and linear regression to learn the relationship between the likelihood function and AHRM parameters, which took about 30 minutes. The differentiable functional relationship enabled efficient exploration of a large parameter space in fitting the models (<1 ms per sample).

In this paper, we provide an overview of the AHRM as a generative model of trial-by-trial human performance in perceptual learning. We also introduce a Bayesian inference procedure (BIP) used to independently estimate the posterior distribution of AHRM parameters for each subject. Subsequently, we present the HB-AHRM, designed to collectively estimate the joint posterior distribution of hyperparameters and parameters at multiple levels of the hierarchy. We discuss the simulation technologies, including PyTensor, and the method for likelihood function approximation. These procedures are applied to data from Petrov et al. (2005). Our analysis involves comparing the goodness of fit the BIP and HB-AHRM, and evaluating the variability of estimated AHRM parameters, learning curves and weight structures. In addition, we conducted a simulation study to evaluate parameter recovery and HB-AHRM’s ability in predicting the performance of a new simulated observer with no or limited training data.

## THEORETICAL DEVELOPMENT

### The Augmented Hebbian Reweighting Model (AHRM)

In this section, we briefly describe the augmented Hebbian reweighting model (AHRM). More details of the model can be found in the original paper (Petrov et al., 2005).

#### Representation module

For subject *i* in test *j*, the stimulus image consists of a signal image *S_ijt_* and an external noise image *N_ijt_* in each trial *t*. The representation module encodes the stimulus image into expected activations over a bank of orientation and spatial-frequency channels tuned to different orientations *φ* and spatial frequencies *f*, *E*5*φ*, *f*7*S_ijt_*, *N_ijt_*8, through convolution, halfwave rectification, contrast normalization and pooling over phase and space (Petrov et al., 2005). We consider 35 *channels* centered at 7 orientations (*φ*ɛ[0°, ±15°, ±30°, ±45°]) and 5 spatial frequencies (*f* ɛ[1, 1.4, 2, 2.8, 4] cycles per degree). The expected activation is then combined with internal noise *ε_φ,f_* (with mean 0 and standard deviation σ*_r_*) and passed to a saturating non-linearity to compute the activation in each channel:

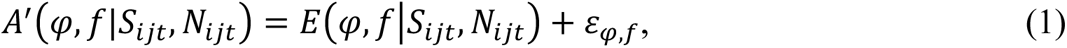

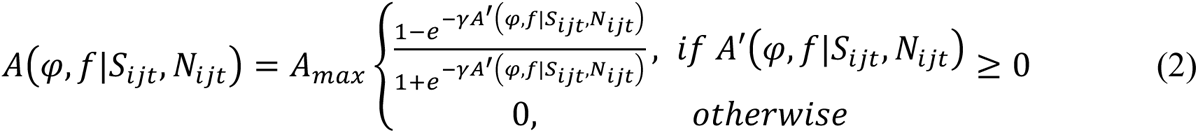

#### Task-specific decision module

The decision module weighs the evidence in the noisy activations from the representation module to generate a response in each trial. Specifically, it first aggregates the activation pattern *A*(*φ*, *f*|*S_ijt_*, *N_ijt_*) over the orientation and spatial-frequency channels using current weights *w_φ,f_*(*t*), a top-down bias *b(t),* and a Gaussian decision noise *ε* (mean 0 and standard deviation *σ_d_*):

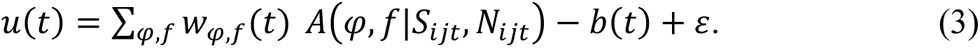

and then computes its output *o*(*t*) as a sigmoidal function of *u(t)*:

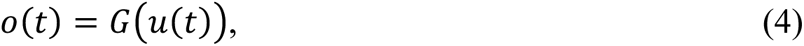

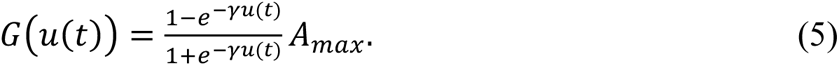

The decision variable *o*(*t*) saturates at ±*A_max_* = ±0.5; the model responds left or counterclockwise if *o*(*t*) < 0, and right or clockwise otherwise.

The weights are initiated to be proportional to the preferred orientation of the representation module relative to the vertical: *w_φ,f_*(0) = (*φ*/30°)*w_init_*.

#### Learning module

The weights between the representation and decision modules are updated on each trial using an augmented Hebbian reweighting rule, in which feedback, *F*(*t*) = ± 0.5, is used as the correct output of the decision module. The change of weight *w_φ,f_*(*t*) in each channel depends on the activation of the representation *A*(*φ*, *f*|*S_ijt_*, *N_ijt_*), the correct output of the decision module and the internal learning rate *η*:

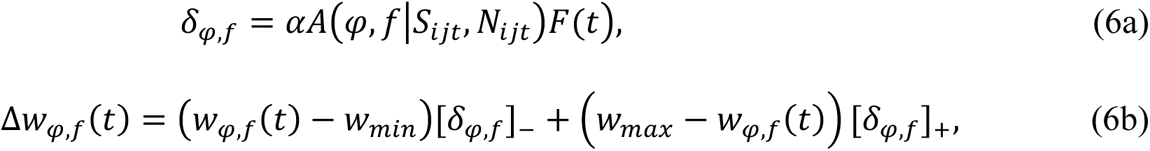

where *w_min_* = −1 and *w_max_* = 1.0.

#### Adaptive criterion control

The adaptive criterion control module shifts the decision variable on each trial to compensate for biases in the immediate history of responses by adding a bias correcting term to the activation at the decision module. It assumes that observers monitor their own responses and seek to equalize response frequencies—trying to match stimulus probabilities that are often balanced in experiments. A weighted running average of recent responses exponentially discounts the distant past response history:

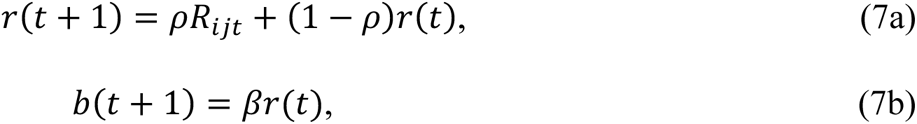

where R(t) is the response in the current trial, and r(t) is the weighted running average responses, and b(t) is the bias. Following Petrov et al. (2005), we set *ρ*=0.02.

In summary, the AHRM has six free parameters (Table 1), internal learning rate *α*, bias strength *β*, activation function gain *γ*, standard deviation of decision noise *σ_d_*, standard deviation of representation noise *σ_r_*, and initial weight scaling factor *w_init_*. Additional parameters, including maximum activation level, weight bounds, orientation spacing, and spatial frequency spacing, are fixed.

**Table 1:** AHRM parameters and their corresponding symbols in the BIP and HB-AHRM.

| Parameters | AHRM | BIP/HB-AHRM | Values |
| --- | --- | --- | --- |
| Learning rate | $\alpha$ | $\theta_{ij1}$ | |
| Bias strength | $\beta$ | $\theta_{ij2}$ | |
| Activation function gain | $\gamma$ | $\theta_{ij3}$ | |
| Decision noise | $\sigma_d$ | $\theta_{ij4}$ | |
| Representation noise | $\sigma_r$ | $\theta_{ij5}$ | |
| Initial weight scaling factor | $w_{init}$ | $\theta_{ij6}$ | |
| Maximum activation level | $A_{max}$ | | 0.5 |
| Weight bounds | $w_{min/max}$ | | $\pm 1$ |
| Orientation spacing | $\Delta\varphi$ | | $15^\circ$ |
| Spatial frequency spacing | $\Delta f$ | | 0.5 oct |

To simplify notations, we use *θ_i,j_* to denote the AHRM parameters for subject *i* in test *j* (see Table 1 for the correspondence with the original AHRM parameters). For a given subject *i* in test *j* with ARHM parameters *θ_i,j_*, we can compute the probability of obtaining a correct response in trial *t*, *p*(*R_ijt_* = 1|θ*_ij_*, *S_ijt_*, *N_ijt_*), and the probability of obtaining an incorrect response in trial *t*, *p*(*R_ijt_* = 0|*θ_ij_*, *S_ijt_*, *N_ijt_*), from repeated simulations of the AHRM. The two probabilities define the likelihood function for each of *T* trials. The likelihood of obtaining all the observed responses of subject *i* in test *j* is the product of all the trial-by-trial likelihoods:

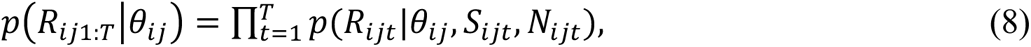

### Bayesian Inference Procedure

The Bayesian Inference Procedure (BIP) is used to estimate the posterior distribution of *θ_ij_* from the trial-by-trial data *Y_ij_* = {(*R_ij_*_1:*T*_, S*_ij_*_1:*T*_, N*_ij_*_1:*T*_,)} of subject *i* in test *j* via Bayes’ rule (Figure 2a):

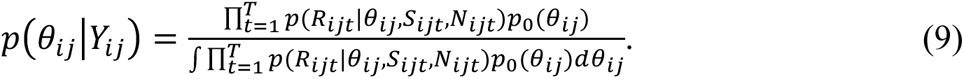

**Figure 2.**
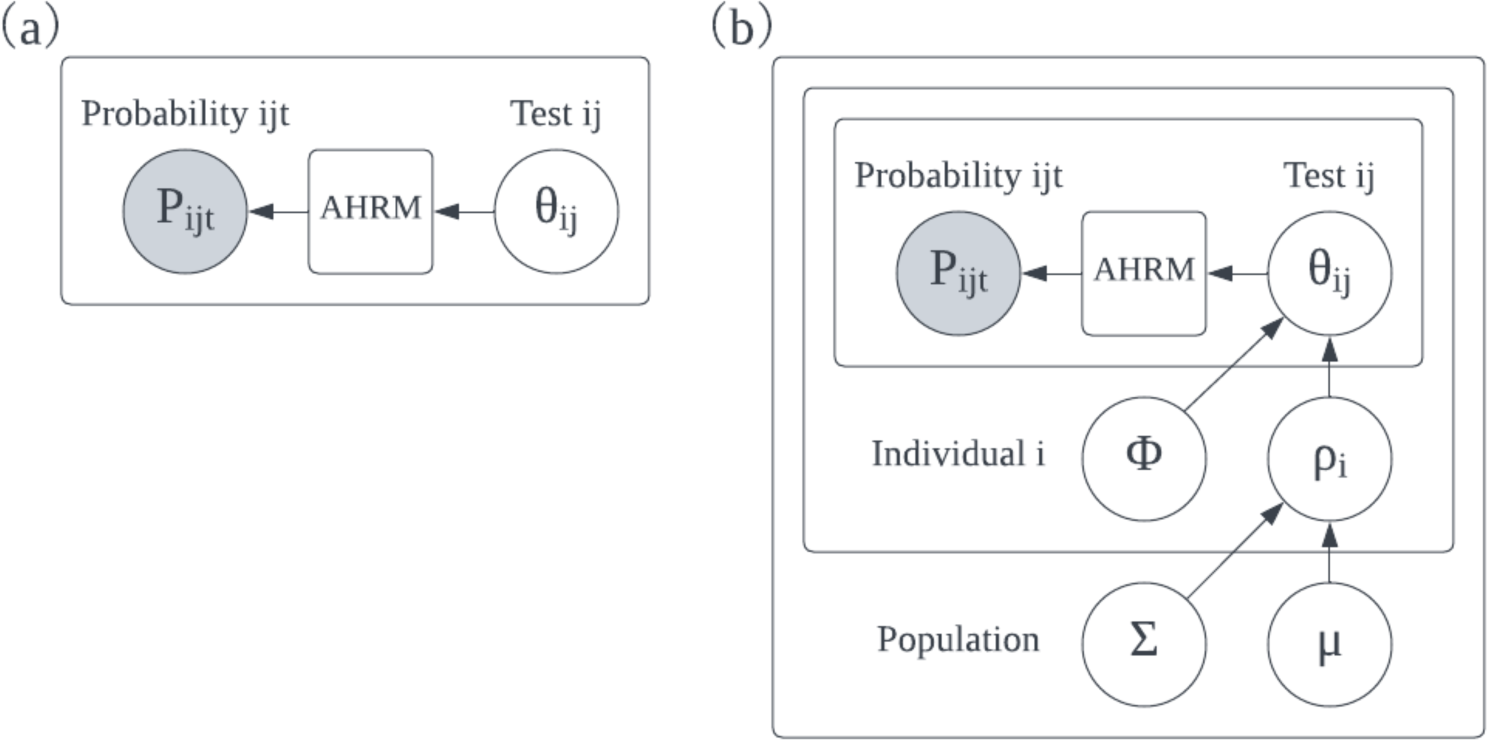
(a) The Bayesian inference procedure (BIP). For a given subject *i* in test *j* with parameters *θ_ij_*, the likelihood of obtaining response *p*(*R_ijt_*) in trial *t* is computed from the AHRM. (b) The HB-AHRM is a three-level hierarchical Bayesian model in which the population level hyperparameter *η* is modeled as a mixture of Gaussian distributions with mean μ and covariance *Σ*, hyperparameter *τ_i_* at the subject level is modeled as a mixture of Gaussian distributions with mean *ρ_i_* and covariance *ϕ*, and the probability distribution of parameters *θ_ij_* is conditioned on *τ_i_*.

Here, *p*(*θ_ij_*|*Y_ij_*) is the posterior distribution of AHRM parameters, *θ_ij_*, given the trial-by-trial data *Y_ij_*, *p*(*R_ijt_*|*θ_ij_*, *S_ijt_*, *N_ijt_*) is the likelihood term, which quantifies the probability of observing responses *R_ijt_* given *θ_ij_*, *S_ijt_*, and *N_ijt_*, *p*_0_(*θ_ij_*) is the prior probability distribution of *θ_ij_*.

The prior of *θ_ij_* is set as a uniform distribution in all its dimension:

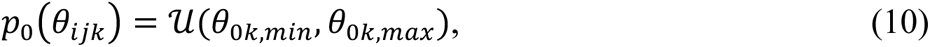

where *θ*_0*k,min*_ and *θ*_0*k,max*_ are the lower and upper bounds of the uniform distribution for dimension *k* (Table 2), which are set based on observed values in prior applications of the model. The denominator of equation (9) is an integral across all possible values of *θ_ij_*.

**Table 2.** Lower and upper bounds of the priors.

| $k$ | 1 | 2 | 3 | 4 | 5 | 6 |
| --- | --- | --- | --- | --- | --- | --- |
| $\theta_{0k,min}$ | 0 | 0 | 0 | 0 | 0 | 0 |
| $\theta_{0k,max}$ | 0.003 | 3 | 3 | 0.3 | 0.1 | 0.3 |

In the BIP, the AHRM parameters are estimated independently for each subject.

### Hierarchical Bayesian Augmented Hebbian Reweighting Model (HB-AHRM)

The HB-AHRM is a three-level hierarchical Bayesian model used to estimate the joint posterior hyperparameter and parameter distribution in all levels, considering covariance within and between subjects (Figure 2b). The HB-AHRM includes probability distributions at the population, subject, and test levels.

#### Population level

The probability distribution of the six-dimensional hyperparameter *η* of the AHRM parameters (Table 1) at the population level is modeled as a mixture of six-dimensional Gaussian distributions with mean μ and covariance *Σ*, which have distributions *p*(*μ*) and *p*(*Σ*):

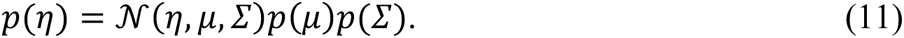

#### Subject level

The probability distribution of hyperparameter *τ_i_* for subject *i* at the subject level is modeled as a mixture of 6-dimensional Gaussian distributions with mean *ρ_i_* and covariance *ϕ*, with distributions *p*(*ρ_i_*|*η*) and *p*(*ϕ*):

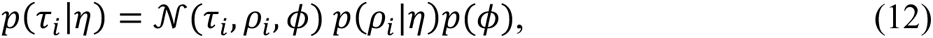

in which *ρ_i_* is conditioned on *η*.

#### Test level

*p*(*θ_ij_*|*τ_i_*), the probability distribution of parameters *θ_ij_* is conditioned on *τ_i_*. The probability of obtaining the entire dataset is computed using probability multiplication, which involves all levels of the model and the likelihood function based on the trial-by-trial data:

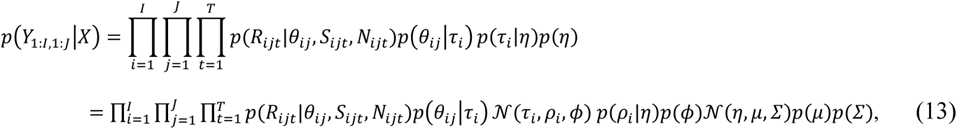

where *X* = (*θ*_1:*I*,1:*J*_, *ρ*_1:*I*_, *μ*, *ϕ*, *Σ*) are all the parameters and hyperparameters in the HB-AHRM.

Bayes’ rule is used to compute the joint posterior distribution of *X*, which includes all HB-AHRM parameters and hyperparameters:

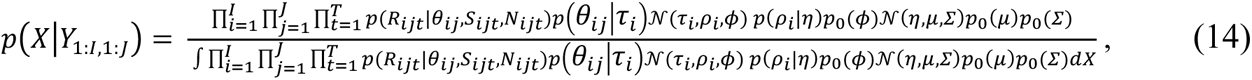

where the denominator is an integral across all possible values of *X* and is a constant for a given dataset and HB-AHRM; *p*_0_(*μ*), *p*_0_(*Σ*), and *p*_0_(*ϕ*) are the prior distributions of *μ*, *Σ*, and *ϕ*, with

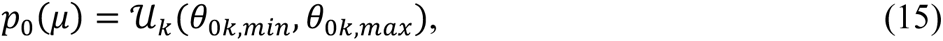

where Ʋ*_k_*(*θ*_0*k,min*_, θ_0*k,max*_) denotes a uniform distribution between *θ*_0*k,min*_ and *θ*_0*k,max*_ in each of the six dimensions, with *θ*_0*k,min*_ and *θ*_0*k,max*_ defined in Table 2. Both *p*_0_(*Σ*) and *p*_0_(*ϕ*) are set with the LKJ distribution with a shape parameter of 2.0 (Lewandowski, Kurowicka, & Joe, 2009).

Equation 14 allows us to estimate the joint posterior distribution of HB-AHRM parameters and hyperparameters across all tests and subjects. Unlike the BIP, the HB-AHRM hyperparameters and parameter estimates mutually constrain each other across tests and subjects via the joint distribution. This allows for more robust and interconnected estimates of HB-AHRM parameters and hyperparameters.

## STUDY 1. RE-ANALYSIS OF Petrov et al. (2005)

### Methods

#### Data

Petrov et al. (2005) investigated perceptual learning in an orientation identification task involving 13 adult subjects with normal or corrected-to-normal vision. Subjects judged the orientation (±10^9^ from vertical) of Gabor patches (windowed sinusoidal gratings, peak spatial frequency=2 c/d) in each trial. The experiment included a nonstationary context where external noise images, predominantly oriented left in context L and right in context R, were superimposed on the target stimuli (Figure 3).

**Figure 3.**
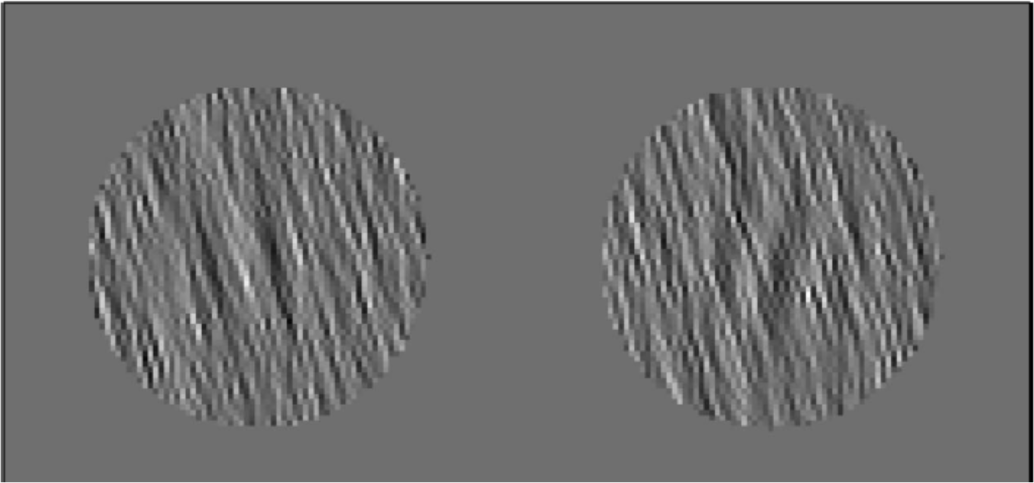
An illustration of left and right titled Gabors in context L.

The study consisted of eight daily sessions, each with four blocks of 300 trials, totaling 9600 trials. Subjects were trained in block sequences of either L-8R-8L-8R-6L-R (7 subjects) or R-8L-8R-8L-6R-L (6 subjects) contexts. Context congruency was randomly selected, with the target Gabor and context either congruent or incongruent in orientation. Gabor contrast was randomly selected from three fixed levels (0.106, 0.160, and 0.245). The resulting behavioral data shows a complex pattern related to congruency and contrast.

The study adhered to ethical standards, with written consent obtained from all subjects prior to the experiment. The research protocol received approval from the institutional review board for human subject research at the University of California, Irvine, and complied with the principles of the Declaration of Helsinki.

#### Likelihood function approximation

Fitting the BIP and HB-AHRM to the data involves using simulations to evaluate the likelihood of a vast set of model parameters. Due to the significant computation time, previous studies relied on grid search methods for AHRM evaluation (Petrov, et al., 2005; 2006). To reduce the computational cost, we approximated the likelihood function by learning its functional relationship with AHRM parameters based on simulated predictions over a large parameter grid. This facilitated fitting the BIP and HB-AHRM models.

We constructed a 6-dimensional mesh grid Θ (Table 3) to train the functional relationship between AHRM parameters and the likelihood function. The mesh grid contained 8 × 8 × 8 × 5 × 5 × 5 = 64,000 sets of model parameters. The ranges and values of the AHRM parameters were chosen based on an exploration of model predictions with various parameters.

**Table 3.** Mesh grid used in the simulation.

| Parameter |  | Values |  |  |  |  |  |  |  |
| --- | --- | --- | --- | --- | --- | --- | --- | --- | --- |
| $\alpha$ | $\theta_{ij1}$ | 0.0001 | 0.0004 | 0.0008 | 0.0012 | 0.0016 | 0.0020 | 0.0024 | 0.0028 |
| $\beta$ | $\theta_{ij2}$ | 0.1 | 0.4 | 0.8 | 1.2 | 1.6 | 2.0 | 2.4 | 2.8 |
| $\gamma$ | $\theta_{ij3}$ | 0.1 | 0.3 | 0.6 | 0.9 | 1.2 | 1.5 | 1.8 | 2.1 |
| $\sigma_d$ | $\theta_{ij4}$ | 0.05 | 0.10 | 0.15 | 0.20 | 0.25 | | | |
| $\sigma_r$ | $\theta_{ij5}$ | 0.025 | 0.050 | 0.075 | 0.100 | 0.125 | | | |
| $w_{init}$ | $\theta_{ij6}$ | 0.05 | 0.10 | 0.15 | 0.20 | 0.25 | | | |

We calculated the likelihood, representing the trial-by-trial probability of a correct response, for each set of AHRM parameters *θ_ij_* across six stimulus conditions over 9600 trials. This computation was based on the average of five simulations, each comprising 300 repeated runs with the same AHRM parameters and a different trial sequence.

The AHRM was used to generate trial-by-trial response based on the set of parameters and the stimulus sequence, using Pytensor library’s *scan* function. Because the exact external noise image on each trial was not available, we obtained a cache of 1200 expected 35-dimensional activations, *E*(*φ*, *f*|*S_ijt_*, *N_ijt_*), consisting of 100 random samples of the 12 combinations of 2 (context) × 3 (Gabor contrast) × 2 (Gabor orientation). In each of the 300 runs, the AHRM starts with the same initial weights and no decision bias, generating orientation judgements in eight sessions with four blocks of 300 trials each. The contexts of the blocks were arranged in terms of L-8R-8L-8R-6L-R or R-8L-8R-8L-6R-L, totaling 9600 trials. Each block of 300 trials consisted of 50 trials in each of the 2 (congruency) × 3 (Gabor contrast) conditions, with a random permutation of the trial types.

Each simulation took 1.6 seconds, in contrast to 22.2 seconds when using a for loop. Additionally, we averaged the likelihoods within each block of 300 trials, resulting in one likelihood for each of the six conditions per block for each set of parameters, and 64,000 likelihoods for the 64,000 sets of AHRM parameters in each of the six conditions per block.

**Figure 4.**
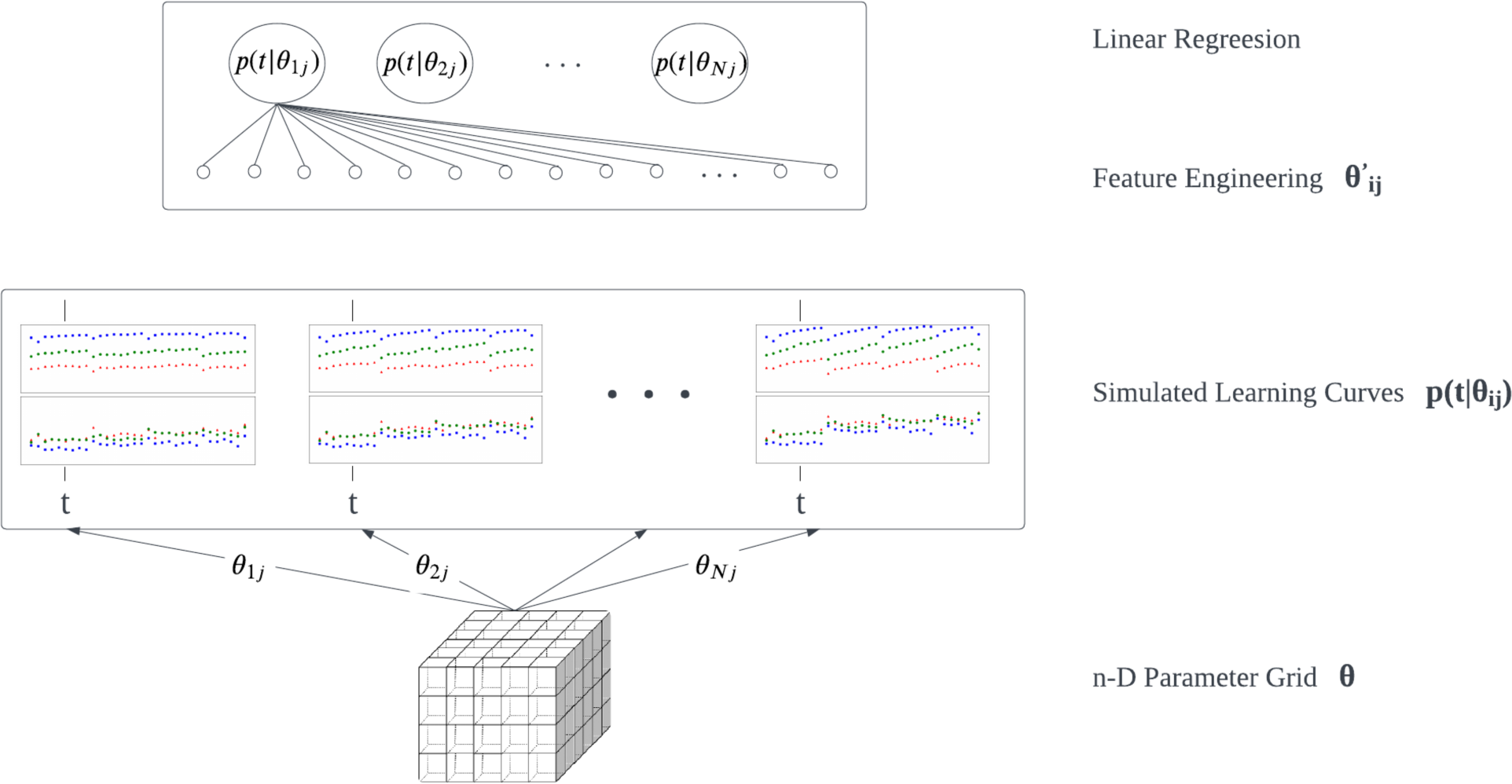
Approximating the likelihood function with feature engineering and linear regression. The simulated learning curves show data for incongruent (top) and congruent (bottom) trials at three contrast levels (colors) over training blocks for each parameter combination.

In each block of the six experimental conditions, we computed the functional relationship between the likelihood and AHRM parameters using feature engineering and linear regression in the Scikit-learn library. Across blocks and experimental conditions, we established a total of 192 functional relationships.

For each relationship, the 64 predictors 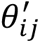 included an intercept, the six AHRM parameters *θ_ij_*, and 57 additional features obtained through comprehensive feature engineering by exploring all 21 quadratic and 36 cubic terms created from the six AHRM parameters. For each block of trials *t*^1^, the linear regression is expressed as:

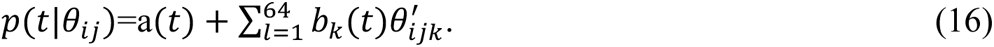

The feature engineering and linear regression step took about half an hour.

The functional relationship for the six conditions in 32 blocks is encoded in a 192 x 64 matrix, where each row represents the coefficients in one experimental condition in one block. This coefficient matrix allows us to calculate the likelihood function *p*(*R_ijt_* = 1|*θ_ij_*, *S_ijt_*, *N_ijt_*) for any AHRM parameters within the mesh grid range, not just the parameter sets in the mesh grid. Moreover, the likelihood functions are differentiable, facilitating various inference functions in *PYMC* (Abril-Pla et al., 2023).

#### Estimating the Posterior Distributions

We utilized the Automatic Differentiation Variational Inference (ADVI) method in the *PYMC* library to generate representative samples of the posterior distributions in the BIP and HB-ARHM. In this method, the variational posterior distribution is assumed to be spherical Gaussian without correlation between parameters and fit to the true posterior distribution. The means and standard deviations of the variational posterior are referred to as variational parameters.

We ran ADVI optimization for 300,000 iterations in the BIP and HB-AHRM to ensure good approximations of the posterior distributions. To generate representative samples of the posterior distribution in the BIP, we used the ADVI method to generate 100,000 samples for each subject *i*. Similarly, we computed 100,000 representative samples of the joint posterior distribution of µ (6 parameters), *Σ* (21 parameters), ρ_ik_ (6 × 13 = 78 parameters), ϕ (21 parameters), and θ_ijk_ (6 × 13 = 78 parameters). A model is considered “converged” when the Evidence Lower Bound (ELBO) stabilizes during iterations, indicating that the variational posterior has adequately approximated the true posterior distribution.

#### Goodness of Fit

We used the Watanabe–Akaike information criterion (WAIC) to compare the BIP and HB-AHRM fits. WAIC quantifies the likelihood of the observed data based on the joint posterior distribution of model parameters while penalizing for model complexity (S. Watanabe & Opper, 2010). Additionally, we assessed the accuracy of model predictions with *RMSE*, the proportion of variance in the observed data explained by the model (R^2^), and the uncertainty of the parameter estimates and model predictions with estimated credible intervals.

The *RMSE* between the predicted and observed quantities is defined as:

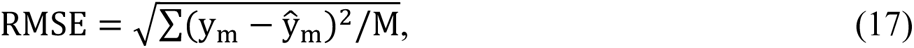

where y_m_ is the observed value, ŷ_m_ is the predicted value, ȳ is the mean of all the observarions, and M is the total number of observations.

The proportion of variance accounted for, R^2^, is defined as:

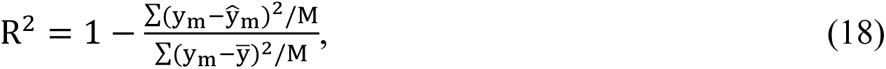

### Results

#### Likelihood function approximation

Figure 5 shows the predicted likelihood function of the AHRM for one set of parameters: *α* = 0.0008, *β* = 1.8, *γ* = 1.2, *σ_d_* = 0.2, σ*_r_* = 0.1, and *w_init_* = 0.17. The model predictions exhibit characteristic patterns observed in Petrov et al. (2005): general learning, persistent switch costs, and within context rapid relearning.

**Figure 5.**
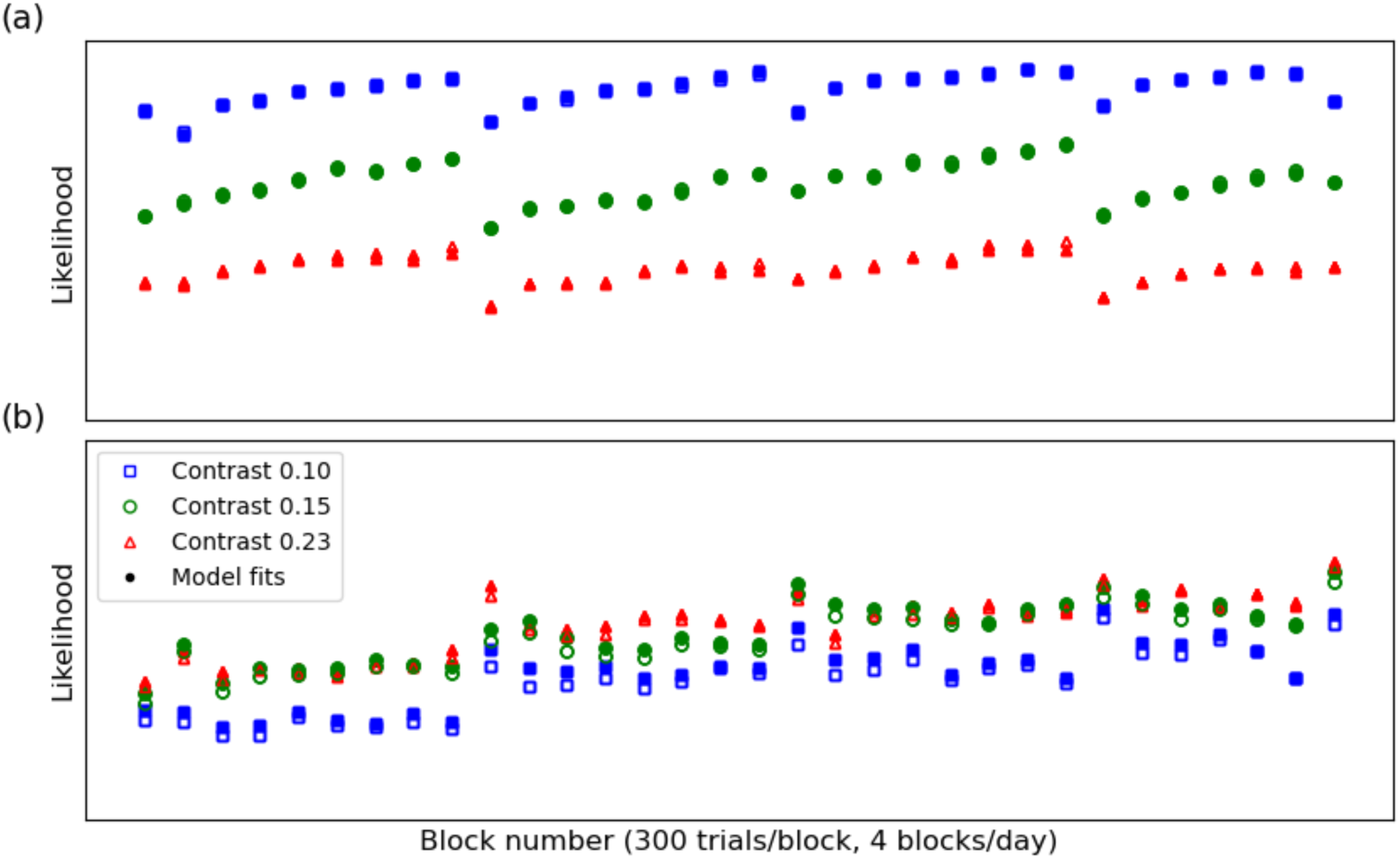
Predicted likelihood function of the AHRM for one set of AHRM parameters (*α* = 0.0008, *β* = 1.8, *γ* = 1.2, *σ_d_* = 0.2, σ*_r_* = 0.1, *w_init_* = 0.17) from the simulations.

Figure 6 shows a scatter plot of the approximate likelihoods from feature engineering and linear regression against likelihoods generated from the AHRM across all 64,000 sets of AHRM parameters in the mesh grid, with an R^2^ of 0.991 and an RMSE of 0.016, indicating excellent approximation of the likelihoods by the linear model (Eq. 16).

**Figure 6.**
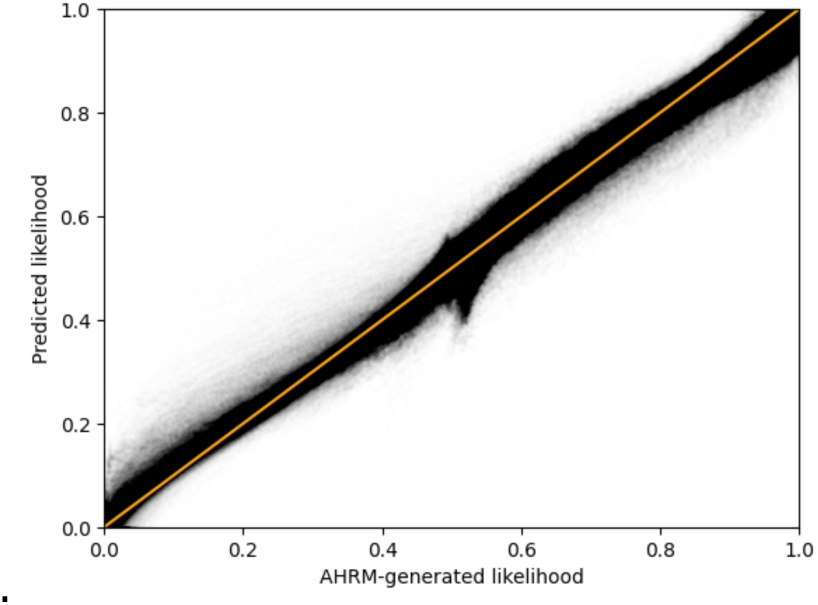
A scatter plot of the approximate likelihoods from feature engineering and linear regression against likelihoods generated from the AHRM.

#### BIP and HB-AHRM Comparison

Both the BIP and HB-AHRM converged, indicated by the stabilization of ELBO.

The WAIC values of the BIP and HB-AHRM were -7908.7 ± 92.5 and -8754.1± 153.3, respectively, with a difference of -845.4 ±179.0. The HB-AHRM provided a significantly better fit to the data. We will focus on the results from the HB-AHRM in the main body of the paper. Detailed results from the BIP can be found in Supplementary Materials A.

#### Posterior Distributions (HB-AHRM)

The marginal posterior distributions of the population-level hyperparameter *η* are depicted in Figure 7. The mean and 94% half width credible interval (HWCI) of these distributions are listed in Table 4. For most *η* components, except *η*_5_ (representation noise), the HWCI was quite small relative to their respective mean.

**Figure 7:**
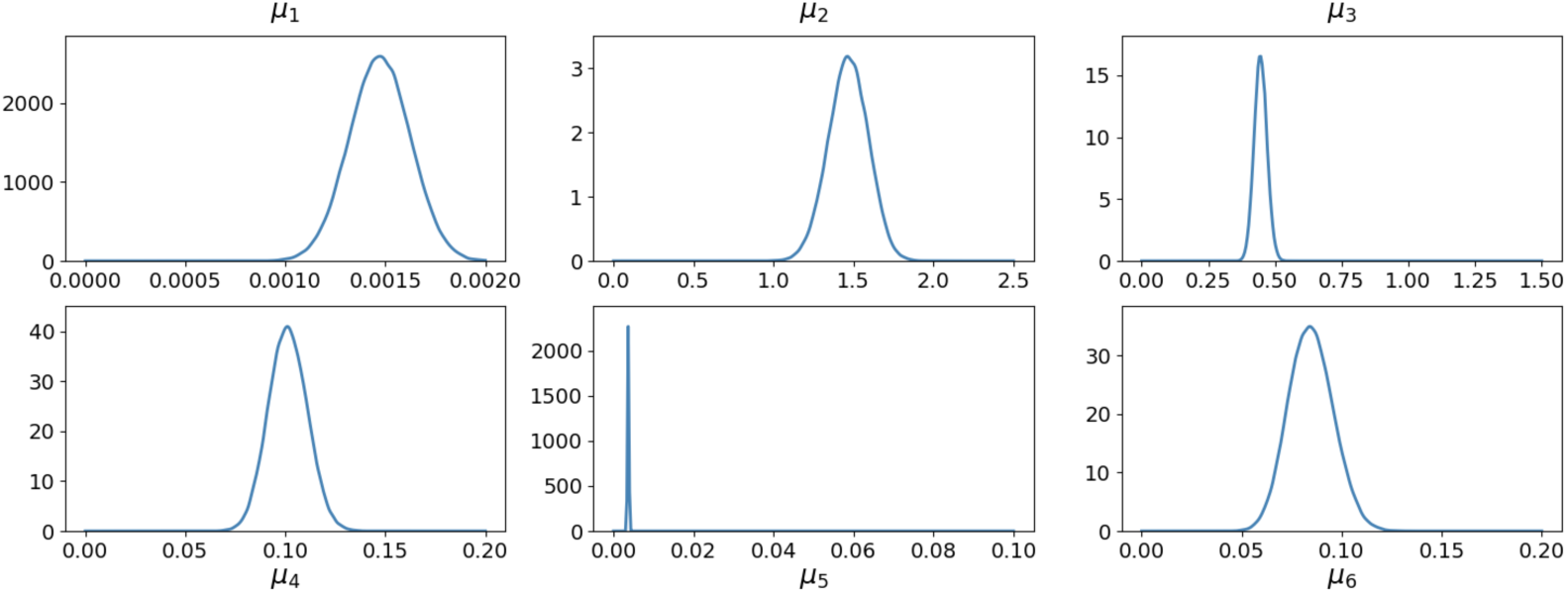
Marginal posterior distributions of the population-level hyperparameter ***η***.

**Table 4.** Mean and 94% HWCI of the marginal distributions of *η*.

| | $\eta_1$ | $\eta_2$ | $\eta_3$ | $\eta_4$ | $\eta_5$ | $\eta_6$ |
| --- | --- | --- | --- | --- | --- | --- |
| Mean | 0.0014 | 1.71 | 0.46 | 0.10 | 0.004 | 0.08 |
| HWCI | 0.0001 | 0.13 | 0.02 | 0.01 | 0.002 | 0.01 |

The expected correlation matrix derived from the expected covariance hyperparameter *Σ* is shown in Table 5. The expected between-subject correlations among *η* components were quite small, and none of them was significantly different from zero, due to the relatively small range of performance variations across the 13 subjects.

**Table 5.** Expected correlation matrix at the population level.

|  |  |  |  |  |  |
| --- | --- | --- | --- | --- | --- |
| 1 | 0.037 | -0.006 | -0.01 | 0.036 | -0.05 |
| 0.037 | 1 | -0.031 | 0.006 | 0.034 | -0.024 |
| -0.006 | -0.031 | 1 | 0.023 | -0.044 | -0.006 |
| -0.01 | 0.006 | 0.023 | 1 | 0.024 | 0.02 |
| 0.036 | 0.034 | -0.044 | 0.024 | 1 | 0.029 |
| -0.05 | -0.024 | -0.006 | 0.02 | 0.029 | 1 |

The marginal posterior distributions of the subject-level hyperparameter τ_i_ for a typical subject (*i* = 6) are depicted in Figure 8. The mean and 94% HWCI of these distributions are listed in Table 6. Compared to *η*, *τ*_6_ components exhibited higher uncertainties, a pattern held across all subjects. The full table with the mean and HWCI of the marginal distributions for all 13 subjects is available in Supplementary Materials B.

**Figure 8:**
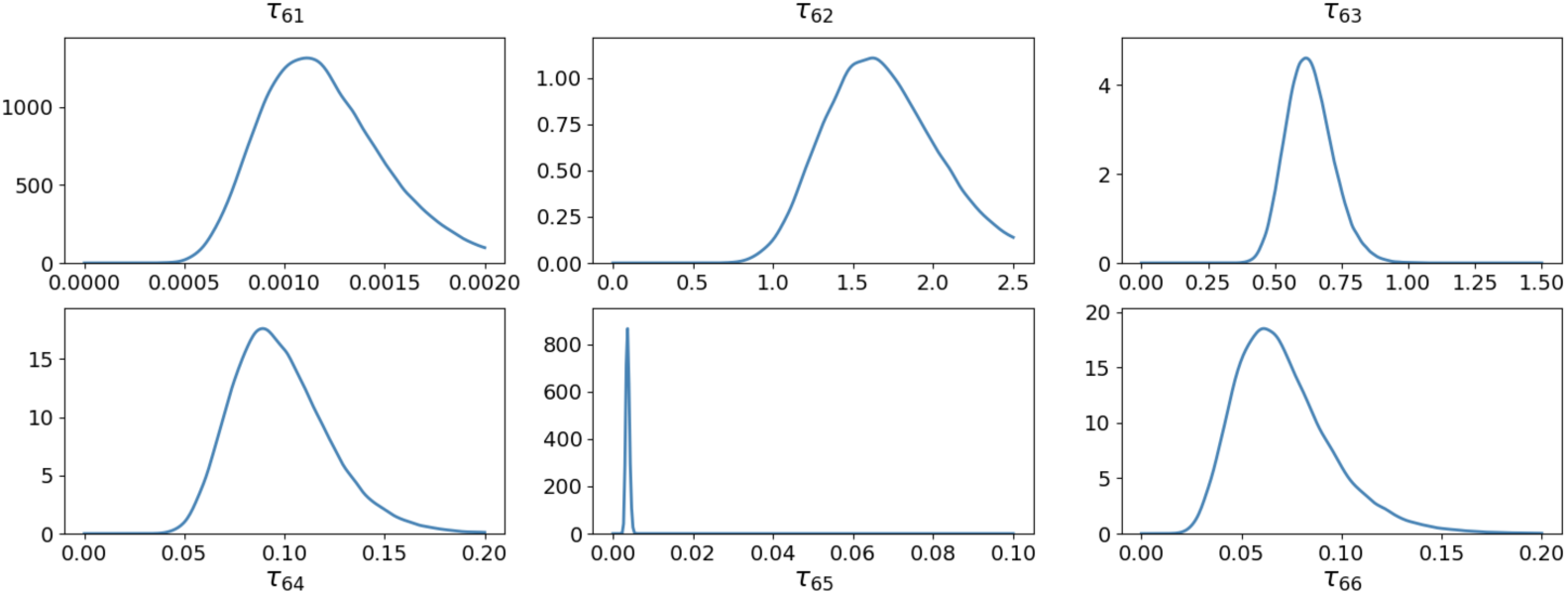
Marginal posterior distributions of the subject-level hyperparameter τ_i_for subject *i* = 6.

**Table 6.**
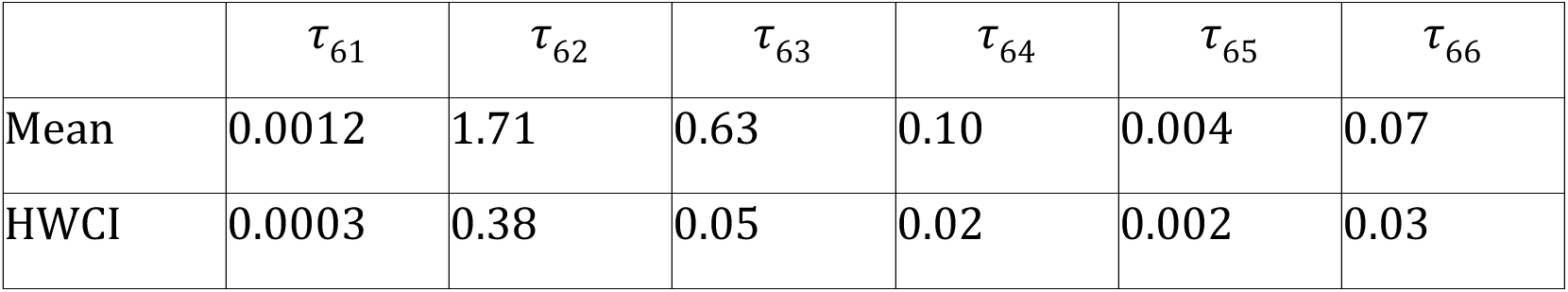
Mean and 94% HWCI of the marginal posterior distributions of τ_6_.

| | $\tau_{61}$ | $\tau_{62}$ | $\tau_{63}$ | $\tau_{64}$ | $\tau_{65}$ | $\tau_{66}$ |
| --- | --- | --- | --- | --- | --- | --- |
| Mean | 0.0012 | 1.71 | 0.63 | 0.10 | 0.004 | 0.07 |
| HWCI | 0.0003 | 0.38 | 0.05 | 0.02 | 0.002 | 0.03 |

The expected correlation matrix derived from the expected covariance hyperparameter Φ is shown in Table 7. The expected correlations between τ_i2_ (bias strength) and τ_i4_ (decision noise) was -0.201 and between components τ_i4_ (decision noise) and τ_i6_ (initial weight scaling factor) was 0.215, both were however not significantly different from zero. The lack of significant within-subject correlations among τ_i_ components suggests that they are essentially independent.

**Table 7.** Expected correlation matrix at the subject level.

|  |  |  |  |  |  |
| --- | --- | --- | --- | --- | --- |
| 1 | -0.085 | 0.075 | 0.093 | -0.008 | 0.052 |
| -0.085 | 1 | 0.074 | -0.201 | -0.029 | 0.023 |
| 0.075 | 0.074 | 1 | -0.014 | 0.006 | -0.095 |
| 0.093 | -0.201 | -0.014 | 1 | 0.004 | 0.215 |
| -0.008 | -0.029 | 0.006 | 0.004 | 1 | -0.004 |
| 0.052 | 0.023 | -0.095 | 0.215 | -0.004 | 1 |

The marginal posterior distributions of the test-level parameter θ_i1_ for subject 6 are depicted in Figure 9. The mean and 94% half width credible interval (HWCI) of these distributions are listed in Table 8. Compared to *η* and τ_6_, θ_61_ components exhibited much lower uncertainties because these test-level parameters are constrained by the experimental data directly. The pattern held across all subjects. The full table with the mean and HWCI of the test-level marginal distributions for all 13 subjects is available in Supplementary Materials B.

**Figure 9.**
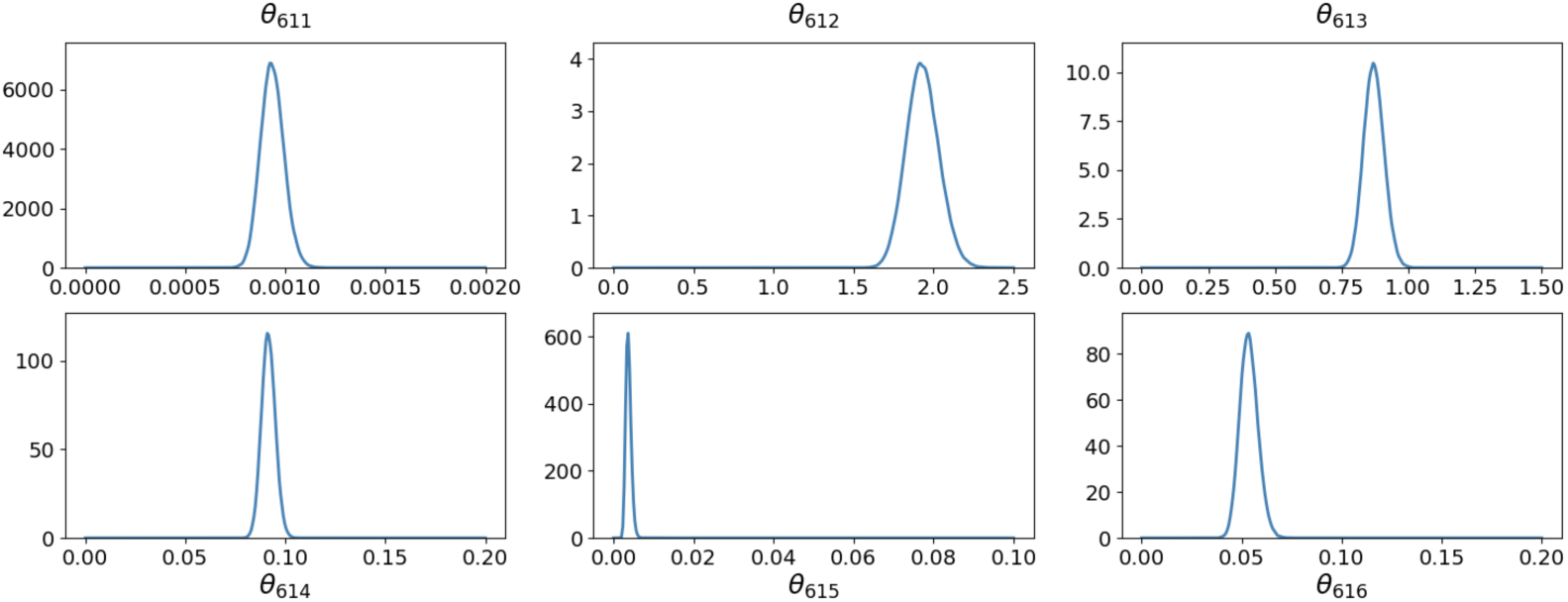
Marginal posterior distributions of the test-level parameter θ_i1_ for subject *i* = 6.

**Table 8.** Mean and 94% HWCI of the marginal posterior distributions of θ_61_.

| | $\theta_{611}$ | $\theta_{612}$ | $\theta_{613}$ | $\theta_{614}$ | $\theta_{615}$ | $\theta_{616}$ |
| --- | --- | --- | --- | --- | --- | --- |
| Mean | 0.0009 | 1.93 | 0.87 | 0.09 | 0.004 | 0.05 |
| HWCI | 0.0001 | 0.10 | 0.04 | 0.00 | 0.001 | 0.00 |

#### Predicted learning curves and weight structures (HB-AHRM)

Figure 10 depicts the observed and predicted population-level z-score learning curves in the incongruent and congruent conditions, as well as the derived d’ learning curves. For the z-score learning curves, the average *RMSE* was 0.173±0.071 and 0.175±0.090 and the average 94% HWCI was 0.064±0.037 and 0.071±0.001 in the congruent and incongruent conditions, respectively, with a *R*^2^ of 0.835.

**Figure 10:**
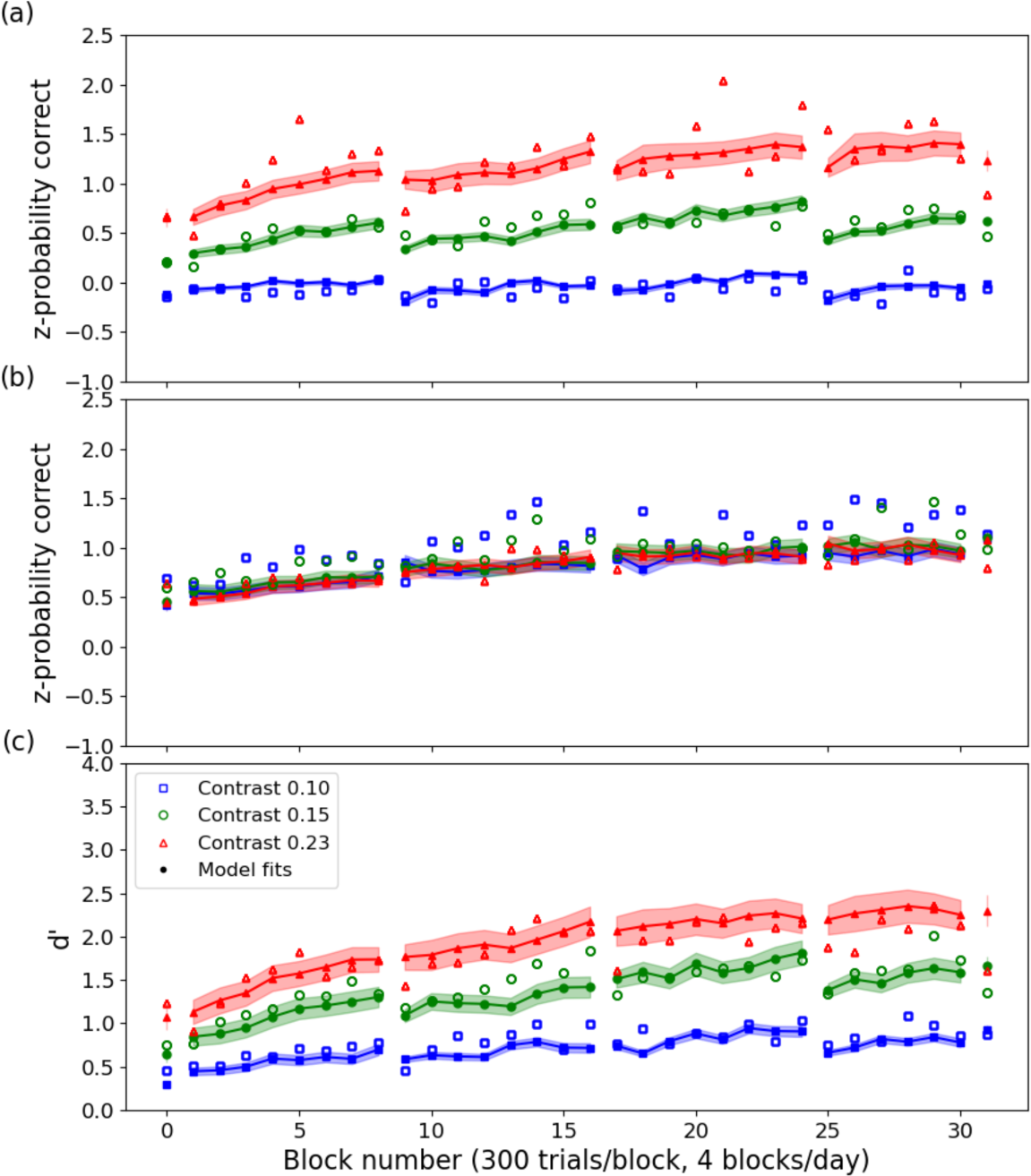
Observed and predicted population-level z-score learning curves in the incongruent (a) and congruent (b) conditions, and the derived *d*’ learning curves (c). Data points represent the average learning curves across all 13 subjects.

For the *d*’ learning curves, the average *RMSE* was 0.199±0.074 and average 94% HWCI was 0.105±0.051, with a *R*^2^ of 0.887. In comparison, the AHRM with parameters from grid search in Petrov et al. (2005) accounted for 88.2% of the variance of the average *d*′ learning curve across all the subject, with a *RMSE* of 0.209. The results suggest that the HB-AHRM provided a largely comparable solution as the original AHRM at the population level. A table that details *RMSE* and 94% HWCI in each of the six experimental conditions is available in Supplementary Materials B.

Figure 11 depicts population-level weight structure evolution over the course of training. Similar to Petrov et al. (2005), the weights for task-relevant channels (e.g., 2 c/d) increased over the course of training, while the weights for task-irrelevant channels (e.g., 4 c/d) stayed more or less the same. The HB-AHRM also allowed us to estimate the uncertainties of the weights. For the channels at 2 c/d, the average HWCI of the weights was 0.008 after 300 trials of training, 0.011 after 2700 trials of training, and 0.011 after 9300 trials of training. For the channels at 4 c/d, the average HWCI of the weights was 0.008 after 300 trials of training, 0.007 after 2700 trials of training, and 0.006 after 9300 trials of training.

**Figure 11:**
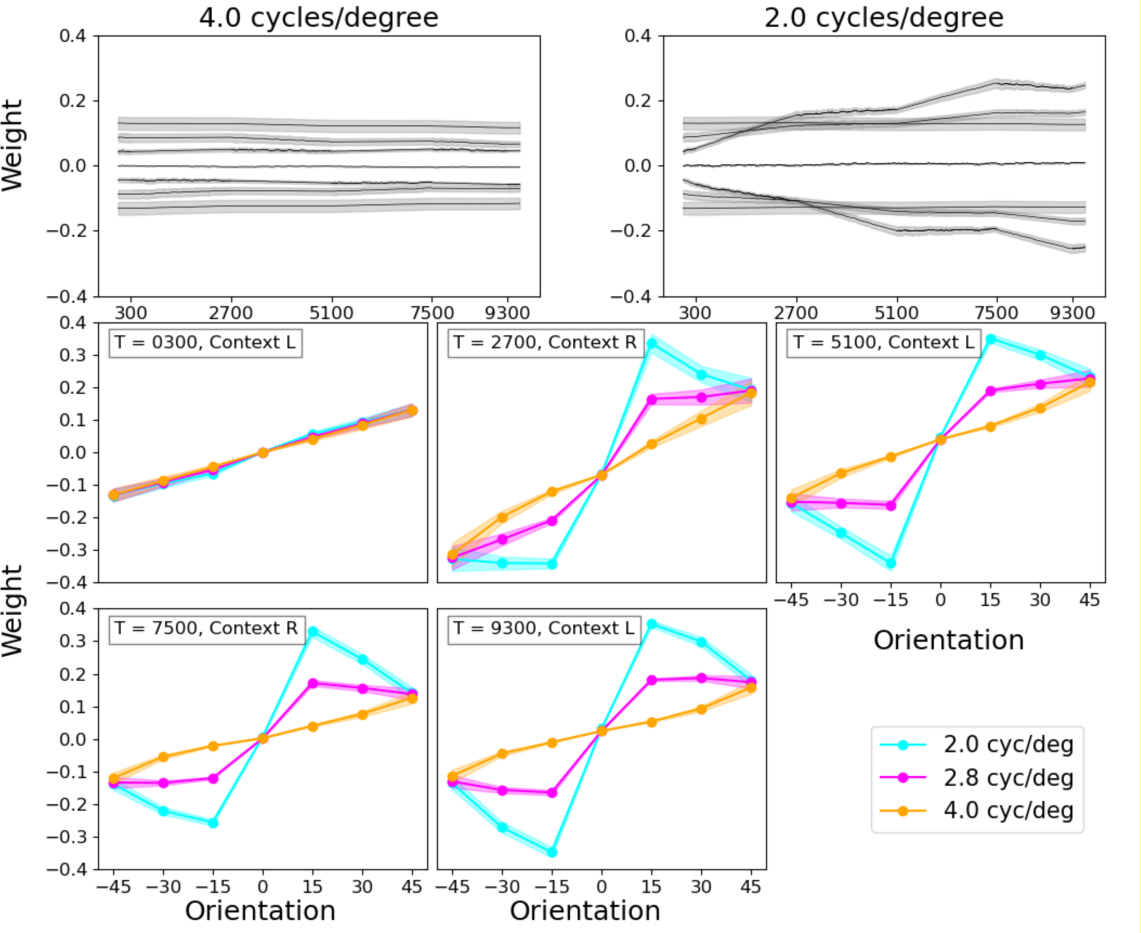
Population-level weight structure evolution over the course of training. Top row: Longitudinal weight traces for units tuned to the target frequency (2.0 c/d) and an irrelevant frequency (4.0 c/d). Each trace corresponds to a particular orientation. Middle and bottom rows: Cross-sections of the weights at 2.0 c/d, 2.8 c/d and 4.0 c/d at the end of each epoch.

Figure 12 depicts the observed and predicted test-level z-score learning curves in the incongruent and congruent conditions, as well as the derived *d*’ learning curves for subject 6. For the z-score learning curves, the average *RMSE* was 0.423±0.204 and 0.456±0.227 and the average 94% HWCI of 0.035±0.001 and 0.032±0.015 in the congruent and incongruent conditions, respectively, with a *R*^2^ of 0.675. For the *d*’ learning curves, the average *RMSE* was 0.276±0.071 and the average 94% HWCI was 0.031±0.014, with a *R*^2^ of 0.744. A table that details *RMSE* and 94% HWCI in each of the six experimental conditions for all the subjects is available in Supplementary Materials B.

**Figure 12:**
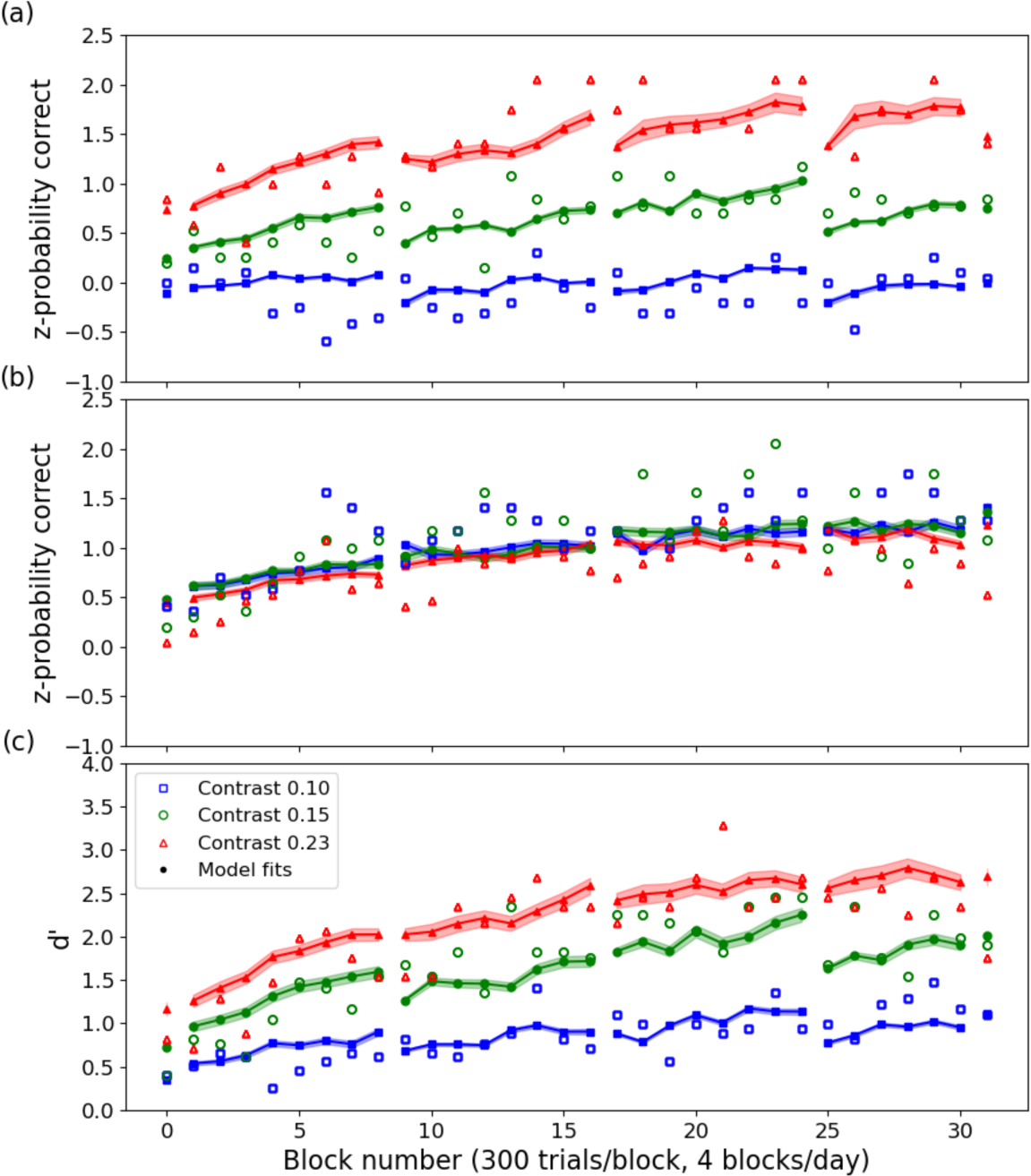
Observed and predicted test-level z-score learning curves in the incongruent (a) and congruent (b) conditions, and the derived d’ learning curves (c) for subject 6.

Figure 13 depicts test-level weight structure evolution over the course of training for subject 6. The pattern of results is very similar to what happened at the population level, although the quantitative weights were different. For the channels at 2 c/d, the average HWCI of the weights was 0.003 after 300 trials of training, 0.005 after 2700 trials of training, and 0.006 after 9300 trials of training. For the channels at 4 c/d, the average HWCI of the weights was 0.003 after 300 trials of training, 0.002 after 2700 trials of training, and 0.001 after 9300 trials of training.

**Figure 13:**
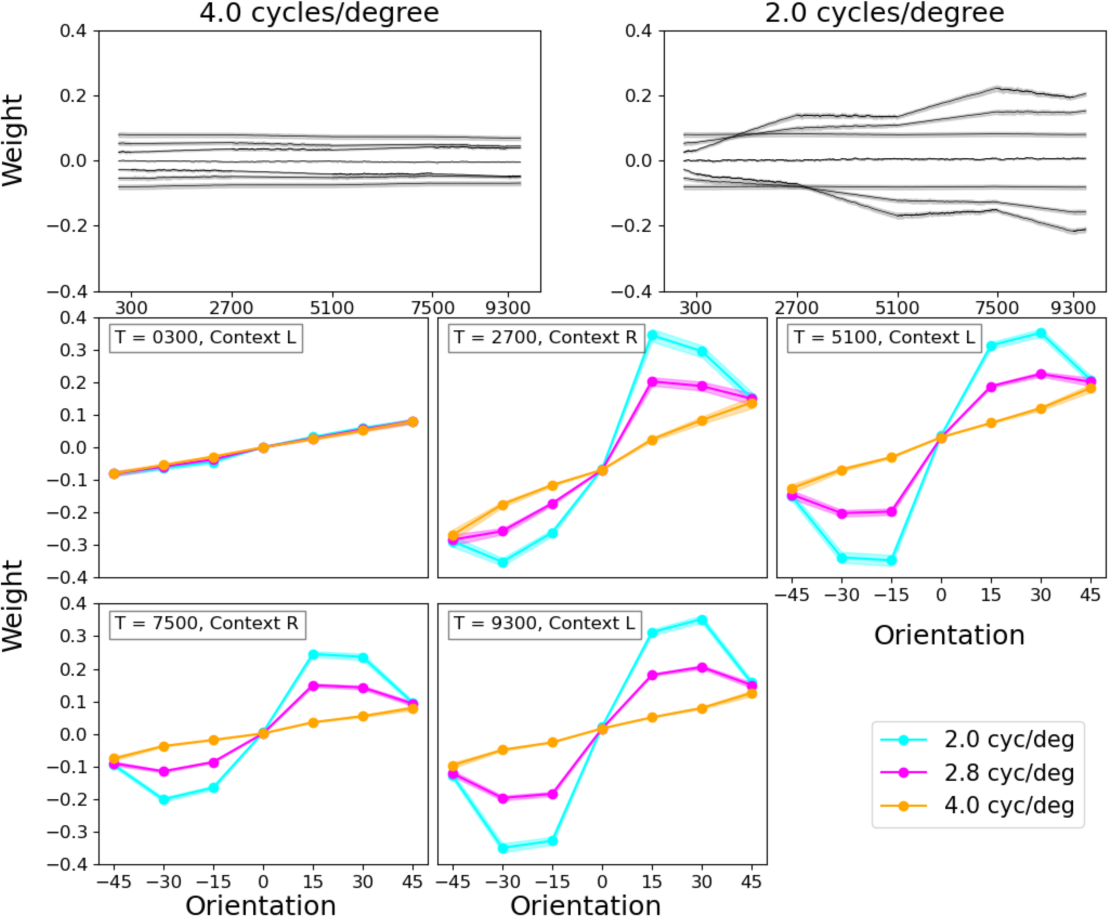
Test-level weight structure evolution over the course of training for subject 6. Top row: Longitudinal weight traces for units tuned to the target frequency (2.0 c/d) and an irrelevant frequency (4.0 c/d). Each trace corresponds to a particular orientation. Middle and bottom rows: Cross-sections of the weights at 2.0 c/d, 2.8 c/d and 4.0 c/d at the end of each epoch.

## STUDY 2: SIMULATIONS

In study 2, we conducted a simulation study to evaluate parameter recovery and HB-AHRM’s ability in predicting the performance of a new simulated observer with no or limited training data.

### Methods

#### Simulated Datasets

We created simulated dataset1 with 13 simulated observers to evaluated parameter recovery. For each simulated observer *i*, we set their AHRM parameters by drawing a random sample from the six-dimensional test-level HB-AHRM posterior distribution θ_i1_ (see Supplementary Materials C for a table of all the parameters). We then created a single trial sequence with 9600 trials based on Petrov et al. (2005) and simulated these observers’ performance using the AHRM with the same trial sequence 300 times. Simulated dataset1 therefore consisted of performance in six experimental conditions, averaged every 300 trials, from each of the 13 simulated observers from the 300 repeated simulations. The structure of the data was identical to that in Petrov et al. (2005).

To evaluate HB-AHRM’s ability in predicting the performance of a new simulated observer with no or limited training data, we created three additional simulated datasets by deleting some training data for a randomly selected subject 13 in simulated dataset1, while keeping all the data from the other 12 simulated observers. Specifically, simulated dataset2, simulated dataset3, and simulated dataset4 comprised 9600 trials of subjects 1-12 and 0 trials, the first 300 trials, and the first 2700 trials of training data for subject 13, respectively.

#### HB-AHRM fitting; statistical evaluation

We fit the HB-AHRM to each of the four simulated datasets and computed the predicted learning curves from the joint posterior distributions of the HB-AHRM hyperparameters and parameters. The procedure was identical to that of STUDY 1.

For simulated dataset1, we evaluated parameter recovery by comparing the mean of the posterior distributions of the AHRM parameters with those of the simulated observers (“the truth”) and computed the 94% HWCI for each of the parameters. We also evaluated the goodness of fit using *RMSE* and *R*^2^.

For simulated dataset2, dataset3, and dataset4, we focused on the predicted learning curves of simulated observer 13 and compared them with the simulated learning curves of the same observer in simulated dataset1 (“the truth”).

### Results

#### Model recovery

As shown in Figures 14 and 15, the HB-ARHM provided excellent fits to simulated dataset1 at both the population and test levels.

**Figure 14:**
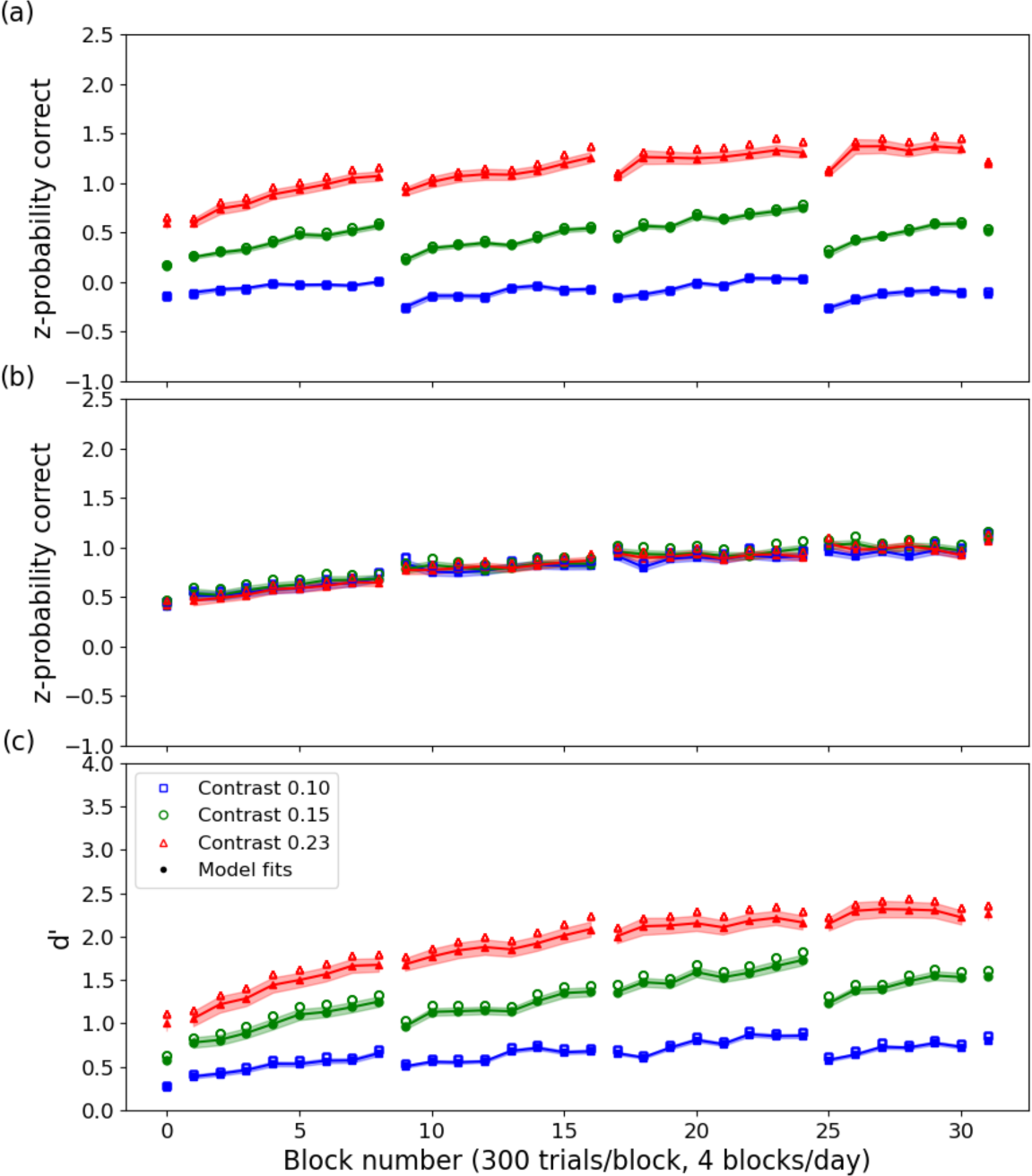
Simulated and predicted population-level z-score learning curves in the incongruent (a) and congruent (b) conditions, and the derived d’ learning curves (c). Data points represent the average simulated learning curves across all 13 simulated observers.

**Figure 15.**
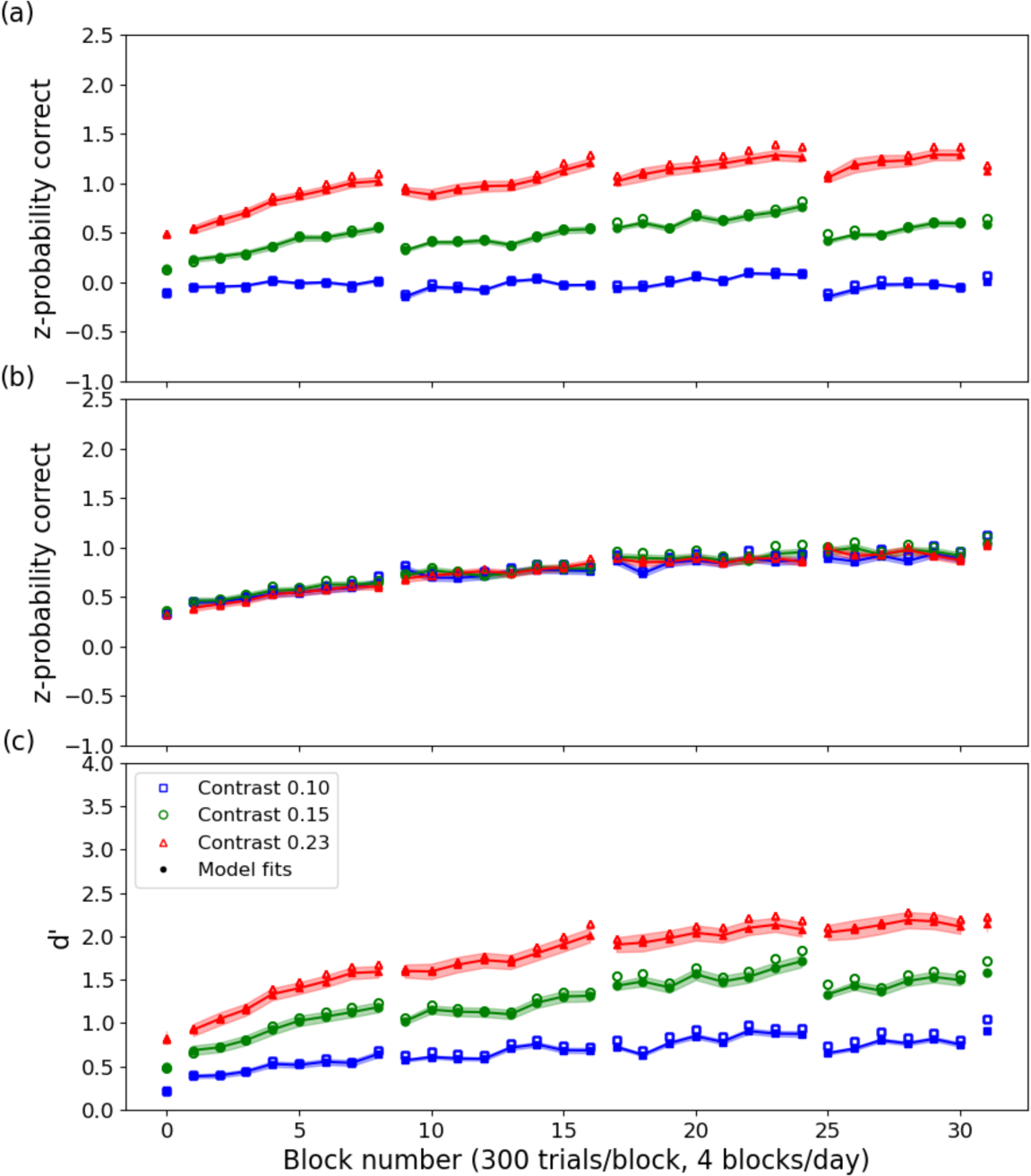
Simulated and predicted test-level z-score learning curves in the incongruent (a) and congruent (b) conditions, and the derived d’ learning curves (c) for simulated observer 13. Data points represent the simulated learning curves.

At the population level, for the z-score learning curves, the average RMSE was 0.014±0.001 and 0.005±0.002 and the average 94% HWCI was 0.037±0.001 and 0.031±0.015, in the congruent and incongruent conditions, respectively, with a *R*^2^ of 0.994. For the *d*’ learning curves, the average RMSE was 0.078±0.038 and the average 94% HWCI was 0.052±0.031, with a *R*^2^ of 0.981. A table that details *RMSE* and 94% HWCI in each of the six experimental conditions is available in Supplementary Materials C.

At the test level, for the z-score learning curves, the average RMSE was 0.010±0.000 and 0.010±0.000 and the average 94% HWCI was 0.034±0.001 and 0.030±0.018 for simulated observer 13 in the congruent and incongruent conditions, respectively, with a *R*^2^ of 0.994. For the *d*’ learning curves, the average RMSE was 0.068±0.023 and the average 94% HWCI was 0.048±0.028, with a *R*^2^ of 0.987. A table that details *RMSE* and 94% HWCI in each of the six experimental conditions for all 13 observers is available in Supplementary Materials C.

Figure 16 shows scatter plots of the recovered AHRM parameters against their true values in simulated dataset1. The *RMSE* between the recovered and true AHRM parameters were 0.00019, 0.308, 0.061, 0.017, 0.002, and 0.011 for the learning rate (*α*), bias strength (*β)*, activation function gain (*γ*), decision noise (*σ_d_*), representation noise (*σ_r_*), and initial weight scaling factor (*w_init_*), respectively. The 94% HWCI for the recovered parameters were 9e-5±2e-5, 0.091±0.040, 0.012±0.002, 0.005±0.001, 7e-17±2e-17, and 0.007±0.001, respectively. For the learning rate, bias strength, decision noise, and initial weight scaling factor, the recovered parameters exhibited excellent correlations with their true values, with Pearson’s correlation coefficients of 0.690, 0.956, 0.877 and 0.980, respectively. For activation function gain, the true values ranged from 0.384 to 0.606, but the recovered values all fell within a narrow range between 0.459 and 0.497, suggesting that the model was not very sensitive to activation function gain. For representation noise, the true values were in a very narrow range (0.00366 to 0.00371), and the recovered values were also in a very narrow range (0.00567 to 0.00593), although with a slightly higher mean. This is because representation noise was very small relative to the external noise in this experiment; it didn’t have much impact on model performance. Overall, these results indicate that the HB-ARHM exhibited very good model recovery.

**Figure 16.**
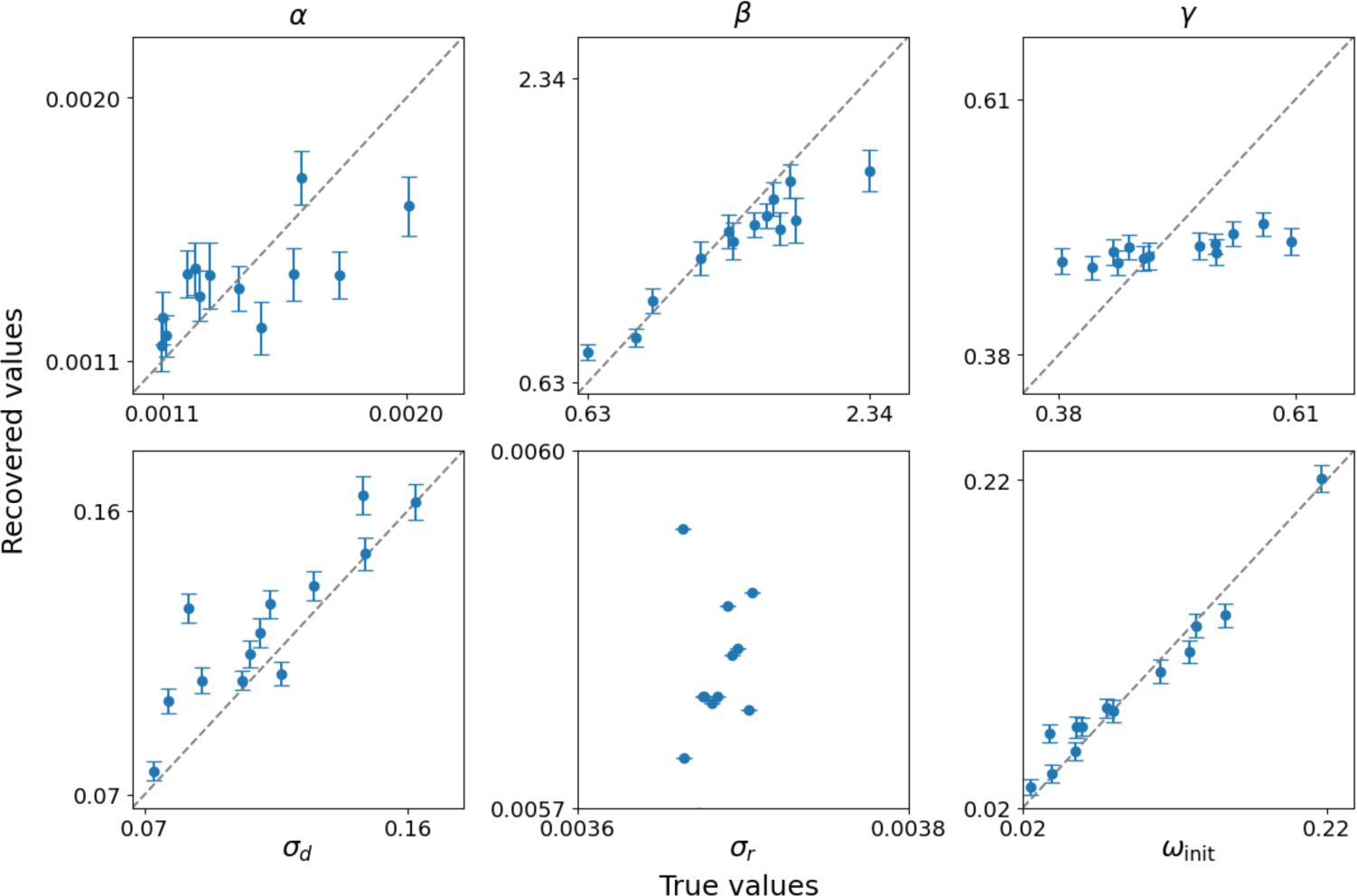
Scatter plots of the recovered AHRM parameters against their true values in simulated dataset1. Each panel shows one AHRM parameter and each point represent one simulated observer. Error bars represent 94% HWCI.

#### Model predictions

Figure 17 shows the predicted learning curves of subject 13 with no data, 30 trials of data, 2700 trials of data, and 9600 trials of data. We compared the predictions with the simulated performance of the subject in simulated dataset1.

**Figure 17.**
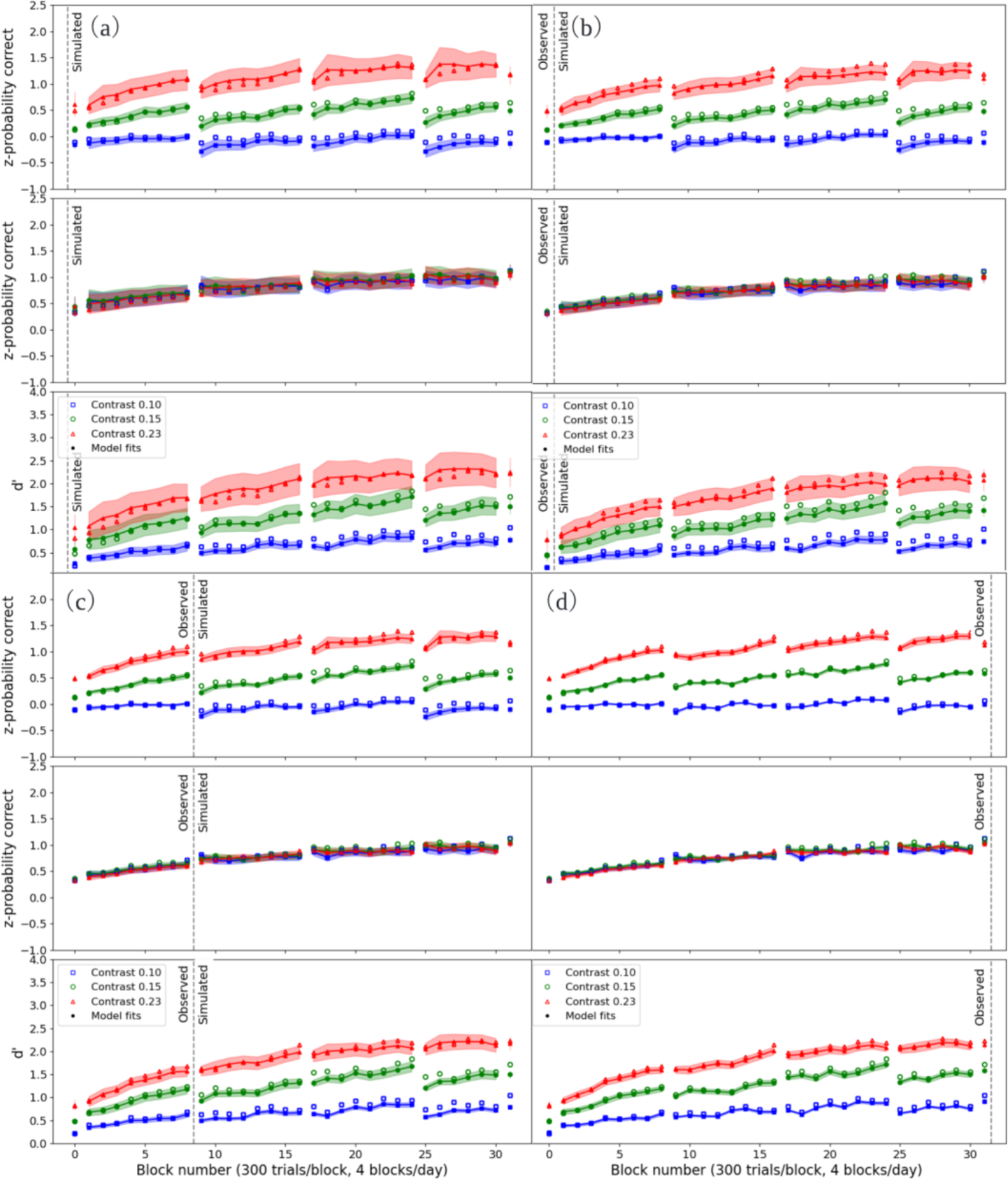
Simulated and predicted z-score and *d*’ learning curves for subject 13 with no data (a), 300 trials of data (b), 2700 trials of data (c), and 9600 trials of data (d). Data points represent the simulated learning curves for the subject in simulated dataset1.

With no data, for the z-score learning curves, the average RMSE was 0.011±0.002 and 0.027±0.009 and the average 94% HWCI was 0.146±0.003 and 0.128±0.068 in the congruent and incongruent conditions, respectively, with a *R*^2^ of 0.972. For the *d*’ learning curves, the average RMSE was 0.104±0.007 and the average 94% HWCI was 0.193±0.115, with a *R*^2^ of 0.960.

With 300 trials of data, for the z-score learning curves, the average RMSE was 0.018±0.006 and 0.025±0.004 and the average 94% HWCI was 0.098±0.002 and 0.088±0.041 in the congruent and incongruent conditions, respectively, with a *R*^2^ of 0.968. For the *d*’ learning curves, the average RMSE was 0.132±0.016 and the average 94% HWCI was 0.146±0.078, with a *R*^2^ of 0.942.

With 2700 trials of data, for the z-score learning curves, the average RMSE was 0.012±0.004 and 0.022±0.006 and the average 94% HWCI was 0.062±0.002 and 0.055±0.022 in the congruent and incongruent conditions, respectively, with a *R*^2^ of 0.980. For the *d*’ learning curves, the average RMSE was 0.101±0.017 and the average 94% HWCI was 0.084±0.044, with a *R*^2^ of 0.966.

Overall, the HB-AHRM made excellent predictions with no data, 300 trials of data, and 2700 trials of data for simulated observer 13. A table that details RMSE and 94% HWCI in each of the six experimental conditions is available in Supplementary Materials C.

## DISCUSSION

In this study, we developed the HB-AHRM and new modeling technologies to address the challenge in fitting the AHRM, a very successful model in visual perceptual learning. A combination of feature engineering and linear regression provided a high-quality approximation of the likelihood function. This approach allowed us to drastically reduce the computation time for fitting the HB-AHRM, estimated to be over 125 days for the dataset in Petrov et al. (2005), which is practically infeasible. The HB-AHRM produced significantly better fits than the BIP, enabling fitting at both the group level with comparable goodness of fit as Petrov et al. (2005), and at the individual level. In stimulation studies, the HB-AHRM along with the new modeling technologies demonstrated robust model recovery properties, accurately predicting simulated observer outcomes with minimal or no initial data.

The AHRM generates trial-by-trial responses based on its parameters and the stimulus sequence. As the exact external noise image for each trial in Petrov et al. (2005) was unavailable, we adopted a procedure similar to the original study, sampling from a cache of 1200 expected activations. The impact of the mismatch with the exact stimulus sequence used in the experiment appeared to be minimal at the group level due to cross-subject averaging, but it may have influenced the fits at the individual subject level because each subject in the original study used a unique random trial sequence. Therefore, it is crucial to accurately record the exact stimulus sequences in future studies.

Many traditional statistical methods assume homogeneity or complete independence across subjects and tests. In contrast, hierarchical modeling (HB) integrates heterogeneous information across multiple levels, using Bayes’ theorem to combine sub-models and probability distributions from all observed data in a study (Kruschke, 2014; Kruschke & Liddell, 2018; Rouder & Lu, 2005). This yields updated joint posterior distributions of hyperparameters and parameters, enhancing accuracy compared to methods that treat each individual independently (H. Gu et al., 2016; Kruschke, 2014). Previous studies, including this one, have shown that HBM provides more precise estimates of parameters of interest compared to traditional methods like the BIP, in estimating contrast sensitivity functions (Zhao, Lesmes, Hou, & Lu, 2021), visual acuity behavioral functions (Zhao, Lesmes, Dorr, & Lu, 2021), and learning curves in perceptual learning (Zhao, Liu, Dosher, & Lu, 2024a, 2024b).

Moreover, HBM offers a robust framework for predictions, treating to-be-predicted performance as missing data to compute their posterior distributions based on available information. Leveraging conditional dependencies across the hierarchy and between- and within-subject covariances, HBM facilitates constructing digital twins and making highly accurate and reasonably precise predictions (Zhao, Lesmes, Dorr, & Lu, 2024).

This study introduces the concept of likelihood function approximation and demonstrates its application in fitting the HB-AHRM. This approach may have broader utility in fitting other stochastic models lacking analytic forms, such as the perceptual template model (Lu & Dosher, 2008), integrated reweighting theory (Dosher et al., 2013), and various response time models (Ratcliff, Smith, Brown, & McKoon, 2016), as well as complex models that requires extensive computations to generate predictions, such as the population receptive field model in retinotopic map studies (Dumoulin & Wandell, 2008). In this particular application, feature engineering and linear regression were employed to generate the approximate likelihood function. Alternatively, other methods, including non-linear regression and machine learning, could be utilized to derive the approximate likelihood function.

In conclusion, we successfully developed and implemented the HB-AHRM using newly developed modeling technologies. These advancements hold promise for widespread applications in fitting stochastic models.

## Supplementary Materials A. BIP Results

The marginal posterior distributions of parameter θ_i1_ for subject 6 from the BIP are depicted in Figure S1. The mean and 94% half width credible interval (HWCI) of these distributions are listed in Table S1.

**Figure S1.**
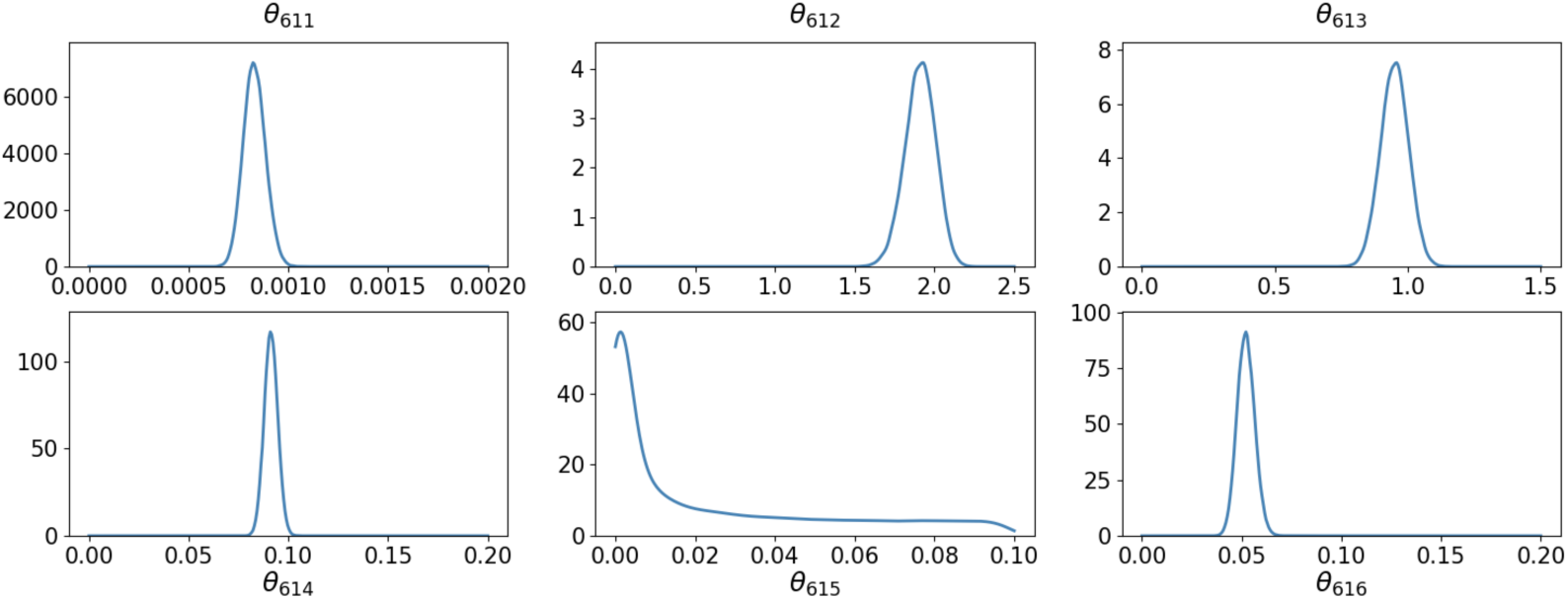
Marginal posterior distributions of parameter θ_i1_ for subject *i* = 6 from the BIP.

**Table S1.** Mean and 94% HWCI of the marginal posterior distributions of θ_i1_.

| | $\theta_{611}$ | $\theta_{612}$ | $\theta_{613}$ | $\theta_{614}$ | $\theta_{615}$ | $\theta_{616}$ |
| --- | --- | --- | --- | --- | --- | --- |
| *Mean* | *0.0008* | *1.91* | *0.95* | *0.09* | *0.054* | *0.05* |
| *HWCI* | *0.0001* | *0.10* | *0.05* | *0.00* | *0.496* | *0.00* |

Figure S2 depicts the observed and predicted test-level z-score learning curves in the incongruent and congruent conditions, as well as the derived *d*’ learning curves from the BIP for subject 6. For the z-score learning curves, the average *RMSE* was 0.403±0.092 and 0.389±0.158 and the average 94% HWCI was 0.039±0.001 and 0.037±0.016 in the congruent and incongruent conditions, respectively, with a *R*^2^ of 0.535±0.090 across all 13 subjects. For the *d*’ learning curves, the average *RMSE* was 0.366±0.057 and the average 94% HWCI was 0.052±0.021, with a *R*^2^ of 0.604±0.135 across all 13 subjects. Tables S2 and S3 detail *RMSE* and 94% HWCI in each of the six experimental conditions.

**Figure S2:**
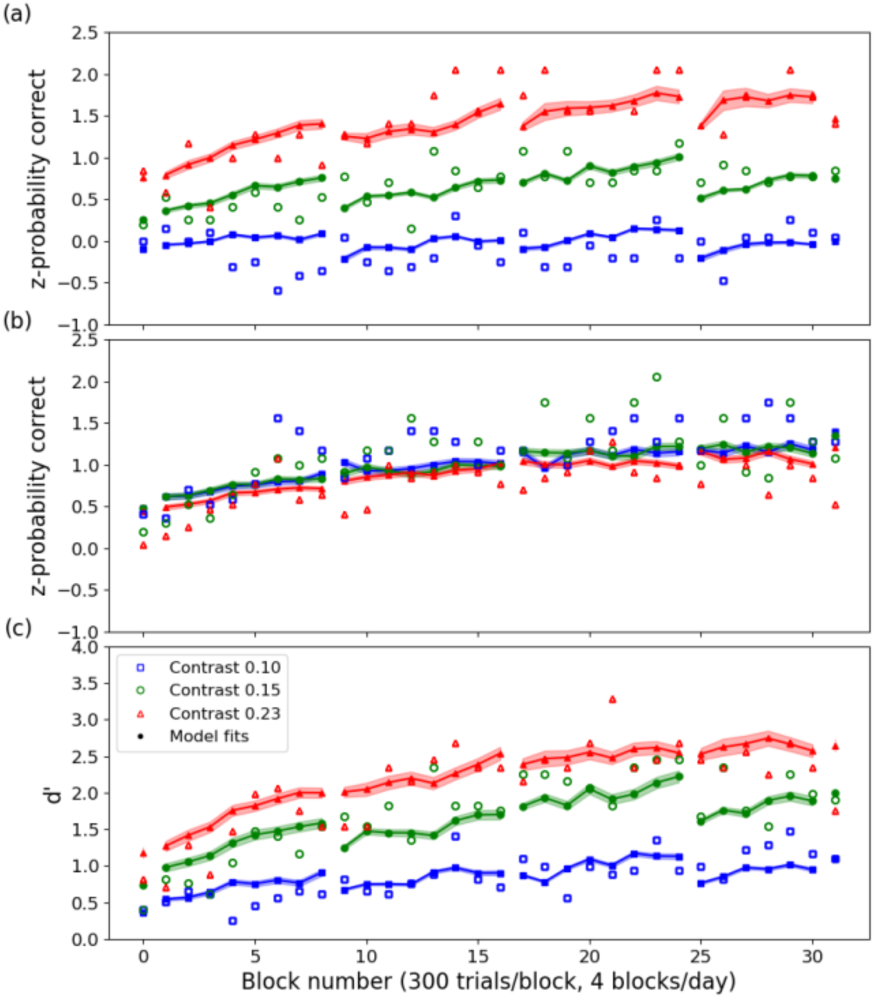
Observed and predicted test-level z-score learning curves in the incongruent (a) and congruent (b) conditions, and the derived d’ learning curves (c) for subject 6.

**Table S2.**
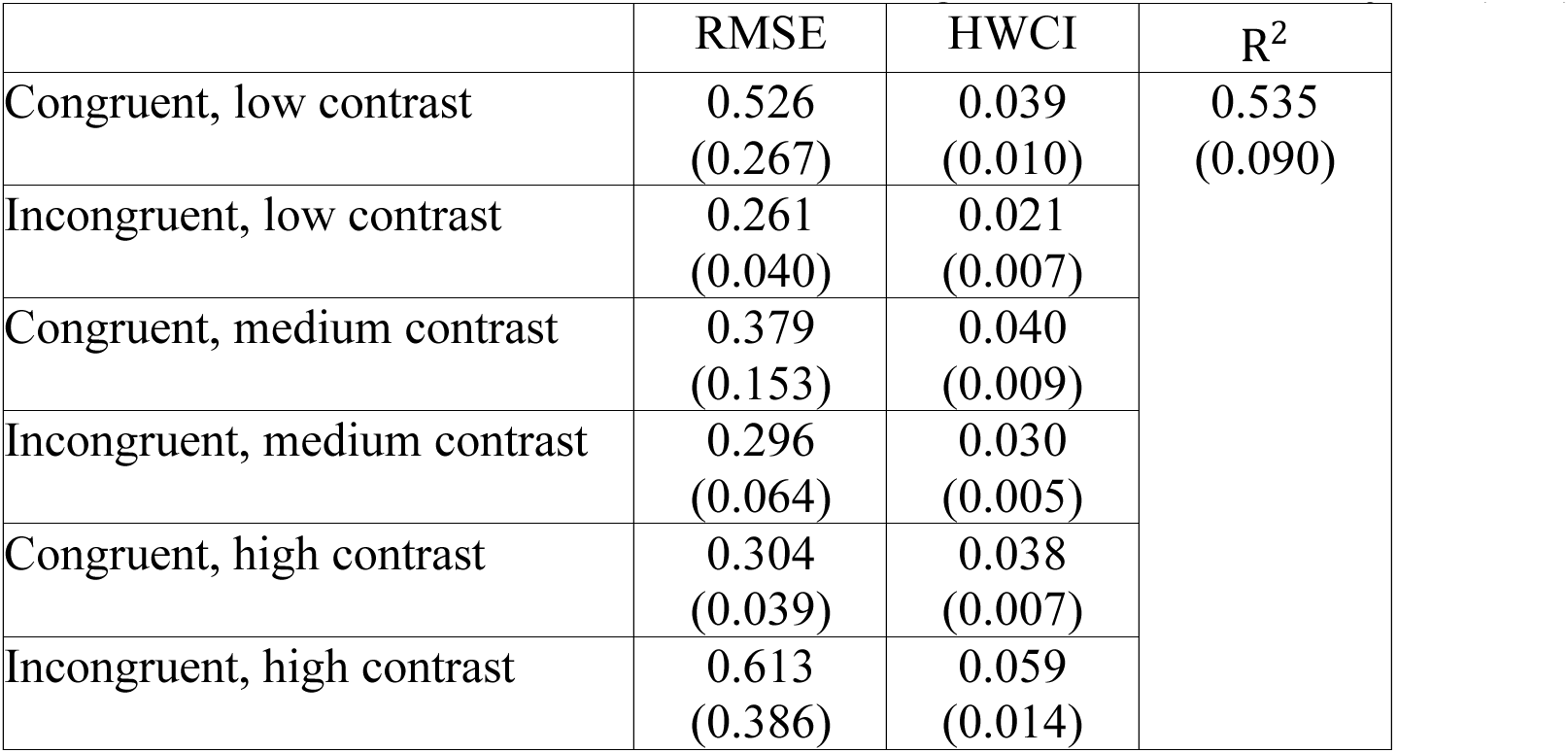
RMSE, 94% HWCI and *R*^2^ for the z-score learning curves across 13 subjects (BIP).

| | RMSE | HWCI | $R^2$ |
| --- | --- | --- | --- |
| Congruent, low contrast | 0.526<br>(0.267) | 0.039<br>(0.010) | 0.535<br>(0.090) |
| Incongruent, low contrast | 0.261<br>(0.040) | 0.021<br>(0.007) |  |
| Congruent, medium contrast | 0.379<br>(0.153) | 0.040<br>(0.009) |  |
| Incongruent, medium contrast | 0.296<br>(0.064) | 0.030<br>(0.005) |  |
| Congruent, high contrast | 0.304<br>(0.039) | 0.038<br>(0.007) |  |
| Incongruent, high contrast | 0.613<br>(0.386) | 0.059<br>(0.014) |  |

**Table S3.** RMSE, 94% HWCI and *R*^2^ for the d’ learning curves across 13 subjects (BIP).

| | RMSE | CI of $d'$ | $R^2$ |
| --- | --- | --- | --- |
| Low contrast | 0.289 | 0.025 | 0.604<br>(0.135) |
| Medium contrast | 0.383 | 0.054 |  |
| High contrast | 0.425 | 0.077 |  |

Figure S3 depicts weight structure evolution over the course of training from the BIP for subject 6. For the channels at 2 c/d, the average HWCI of the weights was 0.003 after 300 trials of training, 0.004 after 2700 trials of training, and 0.006 after 9300 trials of training. For the channels at 4 c/d, the average HWCI of the weights was 0.003 after 300 trials of training, 0.002 after 2700 trials of training, and 0.002 after 9300 trials of training.

**Figure S3.**
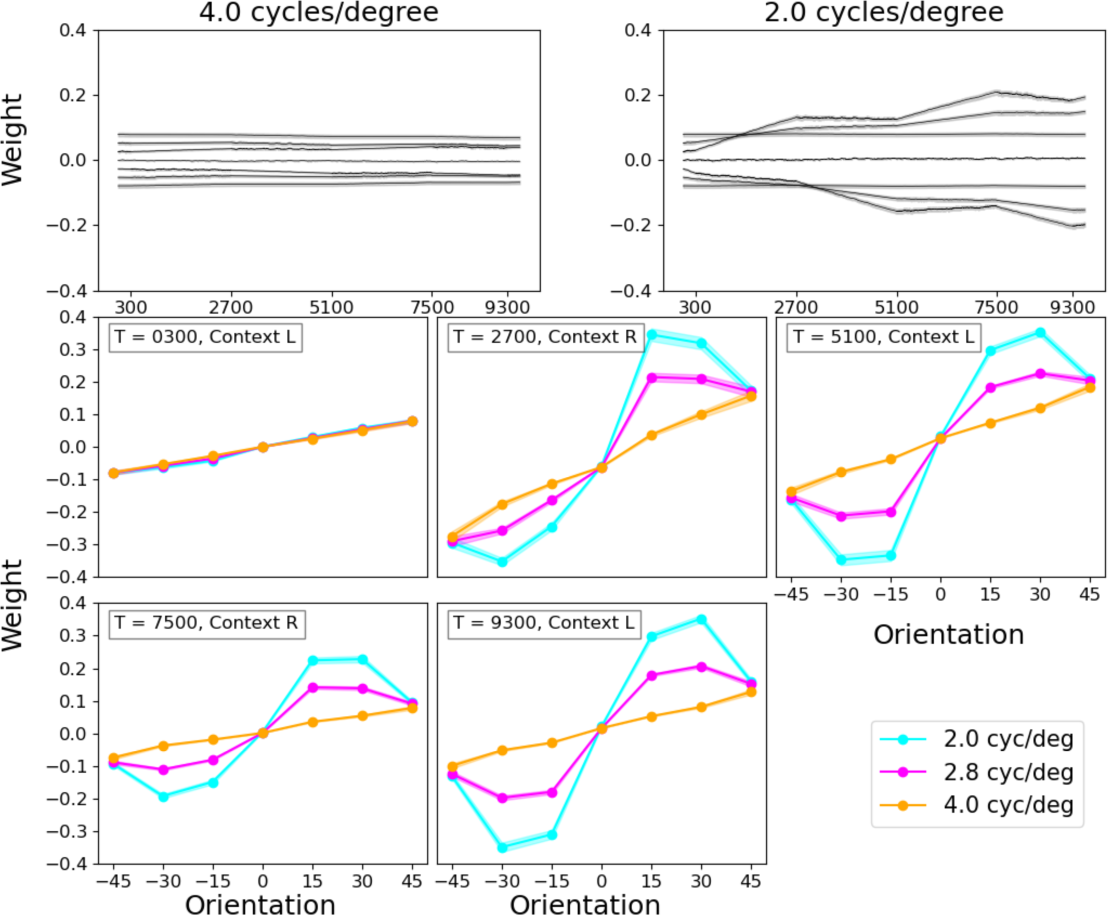
Weight change over time for subject 6 (BIP). Top row: Longitudinal weight traces for units tuned to the target frequency (2.0 c/d) and an irrelevant frequency (4.0 c/d). Each trace corresponds to a particular orientation. Middle and bottom rows: Cross-sections of the weights at 2.0 c/d, 2.8 c/d and 4.0 c/d at the end of each epoch.

### Supplementary Materials B. Additional HB-AHRM Results

**Table S4.** Mean and 94% HWCI of the marginal *τ_ik_* distributions (HB-AHRM).

| $i$ | $\tau_{i1}$ | $\tau_{i2}$ | $\tau_{i3}$ | $\tau_{i4}$ | $\tau_{i5}$ | $\tau_{i6}$ |
| --- | --- | --- | --- | --- | --- | --- |
| **1** | 0.0011 | 1.63 | 0.45 | 0.03 | 0.004 | 0.02 |
|  | (0.0003) | (0.36) | (0.06) | (0.01) | (0.000) | (0.01) |
| **2** | 0.0012 | 1.88 | 0.47 | 0.10 | 0.004 | 0.10 |
|  | (0.0003) | (0.42) | (0.07) | (0.03) | (0.000) | (0.03) |
| **3** | 0.0017 | 1.87 | 0.53 | 0.08 | 0.004 | 0.09 |
|  | (0.0015) | (0.42) | (0.08) | (0.02) | (0.000) | (0.03) |
| **4** | 0.0022 | 1.07 | 0.40 | 0.11 | 0.004 | 0.13 |
|  | (0.0006) | (0.24) | (0.06) | (0.03) | (0.000) | (0.05) |
| **5** | 0.0010 | 1.98 | 0.35 | 0.10 | 0.004 | 0.15 |
|  | (0.0003) | (0.45) | (0.05) | (0.03) | (0.000) | (0.05) |
| **6** | 0.0012 | 1.71 | 0.63 | 0.10 | 0.004 | 0.07 |
|  | (0.0003) | (0.38) | (0.05) | (0.02) | (0.000) | (0.03) |
| **7** | 0.0019 | 0.94 | 0.48 | 0.17 | 0.004 | 0.16 |
|  | (0.0005) | (0.31) | (0.09) | (0.04) | (0.000) | (0.06) |
| **8** | 0.0022 | 1.92 | 0.42 | 0.09 | 0.004 | 0.11 |
|  | (0.0006) | (0.43) | (0.07) | (0.02) | (0.000) | (0.04) |
| **9** | 0.0018 | 0.79 | 0.40 | 0.16 | 0.004 | 0.06 |
|  | (0.0005) | (0.17) | (0.06) | (0.04) | (0.000) | (0.02) |
| **10** | 0.0007 | 1.54 | 0.38 | 0.11 | 0.004 | 0.10 |
|  | (0.0002) | (0.34) | (0.05) | (0.03) | (0.000) | (0.03) |
| **11** | 0.0022 | 1.88 | 0.47 | 0.09 | 0.004 | 0.14 |
|  | (0.0016) | (0.32) | (0.06) | (0.02) | (0.000) | (0.05) |
| **12** | 0.0014 | 1.24 | 0.44 | 0.12 | 0.004 | 0.09 |
|  | (0.0004) | (0.28) | (0.06) | (0.03) | (0.000) | (0.03) |
| **13** | 0.0023 | 1.67 | 0.52 | 0.14 | 0.004 | 0.05 |
|  | (0.0006) | (0.37) | (0.07) | (0.04) | (0.006) | (0.02) |

**Table S5.** Mean and 94% HWCI of the marginal *θ_ij_* distributions (HB-AHRM).

| $i$ | $\theta_{ij}(1)$ | $\theta_{ij}(2)$ | $\theta_{ij}(3)$ | $\theta_{ij}(4)$ | $\theta_{ij}(5)$ | $\theta_{ij}(6)$ |
| --- | --- | --- | --- | --- | --- | --- |
| **1** | 0.0008<br>(0.0001) | 1.74<br>(0.05) | 0.46<br>(0.01) | 0.01<br>(0.00) | 0.004<br>(0.001) | 0.01<br>(0.00) |
| **2** | 0.0010<br>(0.0001) | 2.30<br>(0.13) | 0.49<br>(0.01) | 0.11<br>(0.00) | 0.004<br>(0.001) | 0.11<br>(0.01) |
| **3** | 0.0018<br>(0.0001) | 2.25<br>(0.07) | 0.62<br>(0.01) | 0.07<br>(0.00) | 0.004<br>(0.001) | 0.10<br>(0.01) |
| **4** | 0.0031<br>(0.0003) | 0.77<br>(0.03) | 0.36<br>(0.01) | 0.11<br>(0.00) | 0.004<br>(0.001) | 0.18<br>(0.02) |
| **5** | 0.0006<br>(0.0001) | 2.61<br>(1.14) | 0.27<br>(0.01) | 0.09<br>(0.00) | 0.004<br>(0.091) | 0.24<br>(0.01) |
| **6** | 0.0009<br>(0.0001) | 1.93<br>(0.10) | 0.87<br>(0.04) | 0.09<br>(0.00) | 0.004<br>(0.001) | 0.05<br>(0.00) |
| **7** | 0.0023<br>(0.0002) | 0.59<br>(0.03) | 0.51<br>(0.01) | 0.28<br>(0.01) | 0.004<br>(0.001) | 0.29<br>(0.02) |
| **8** | 0.0030<br>(0.0003) | 2.45<br>(0.11) | 0.37<br>(0.01) | 0.09<br>(0.00) | 0.004<br>(0.001) | 0.13<br>(0.02) |
| **9** | 0.0021<br>(0.0002) | 0.41<br>(0.04) | 0.35<br>(0.01) | 0.23<br>(0.01) | 0.004<br>(0.001) | 0.04<br>(0.01) |
| **10** | 0.0003<br>(0.0001) | 1.61<br>(1.10) | 0.32<br>(0.01) | 0.11<br>(0.00) | 0.004<br>(0.001) | 0.10<br>(0.01) |
| **11** | 0.0031<br>(0.0002) | 2.36<br>(0.06) | 0.48<br>(0.01) | 0.07<br>(0.00) | 0.004<br>(0.001) | 0.20<br>(0.02) |
| **12** | 0.0012<br>(0.0001) | 1.02<br>(0.06) | 0.43<br>(0.01) | 0.14<br>(0.01) | 0.004<br>(0.001) | 0.10<br>(0.01) |
| **13** | 0.0035<br>(0.0002) | 1.84<br>(0.13) | 0.59<br>(0.02) | 0.19<br>(0.01) | 0.004<br>(0.001) | 0.04<br>(0.01) |

**Table S6.**
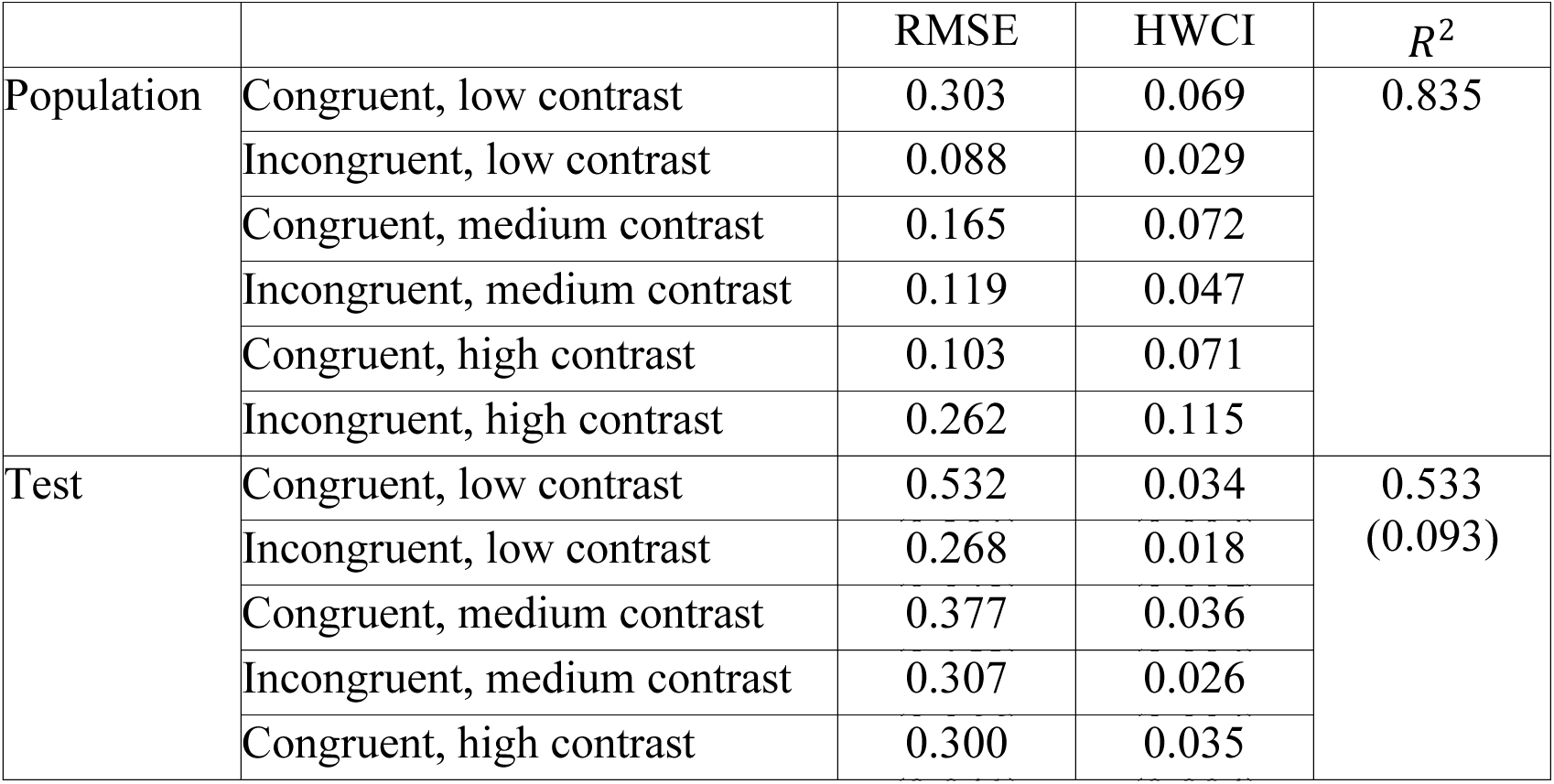
RMSE, HWCI and *R*^2^ for the z-score learning curves at the population and test levels.

**Table S7.** RMSE, HWCI and *R*^2^ for the *d*’ learning curves at the population and test level.

| | | RMSE | HWCI | $R^2$ |
| --- | --- | --- | --- | --- |
| Population | Low contrast | 0.135 | 0.043 | 0.887 |
|  | Medium contrast | 0.157 | 0.103 |  |
|  | High contrast | 0.305 | 0.168 |  |
| Test | Low contrast | 0.287 | 0.023 | 0.615<br>(0.115) |
|  | Medium contrast | 0.303 | 0.050 |  |
|  | High contrast | 0.418 | 0.075 |  |

**Table S8.** Parameters of the Simulated Observers.

| $i$ | $\theta_{i1}(1)$ | $\theta_{i1}(2)$ | $\theta_{i1}(3)$ | $\theta_{i1}(4)$ | $\theta_{i1}(5)$ | $\theta_{i1}(6)$ |
| --- | --- | --- | --- | --- | --- | --- |
| 1 | 0.0011 | 1.48 | 0.41 | 0.11 | 0.004 | 0.15 |
| 2 | 0.0014 | 1.31 | 0.47 | 0.11 | 0.004 | 0.02 |
| 3 | 0.0016 | 1.89 | 0.52 | 0.15 | 0.004 | 0.08 |
| 4 | 0.0015 | 1.02 | 0.38 | 0.14 | 0.004 | 0.13 |
| 5 | 0.0012 | 0.92 | 0.53 | 0.16 | 0.004 | 0.08 |
| 6 | 0.0011 | 1.86 | 0.46 | 0.09 | 0.004 | 0.13 |
| 7 | 0.0020 | 1.79 | 0.55 | 0.12 | 0.004 | 0.11 |
| 8 | 0.0012 | 0.63 | 0.45 | 0.09 | 0.004 | 0.04 |
| 9 | 0.0016 | 1.71 | 0.58 | 0.07 | 0.004 | 0.06 |
| 10 | 0.0011 | 1.75 | 0.44 | 0.08 | 0.004 | 0.05 |
| 11 | 0.0013 | 1.64 | 0.53 | 0.10 | 0.004 | 0.22 |
| 12 | 0.0017 | 1.51 | 0.43 | 0.13 | 0.004 | 0.04 |
| 13 | 0.0013 | 2.34 | 0.61 | 0.11 | 0.004 | 0.06 |

### Supplementary Materials C. Additional Results in the Simulation Study

**Table S9.** Mean and 94% HWCI of the η distributions.

| | $\eta(1)$ | $\eta(2)$ | $\eta(3)$ | $\eta(4)$ | $\eta(5)$ | $\eta(6)$ |
| --- | --- | --- | --- | --- | --- | --- |
| Mean | 0.0014 | 1.37 | 0.48 | 0.12 | 0.006 | 0.08 |
| HWCI | 0.0003 | 0.08 | 0.01 | 0.01 | 0.000 | 0.01 |

**Table S10.** Mean and 94% HWCI of the *τ_ik_* distributions (Simulation study).

| $i$ | $\tau_{i1}$ | $\tau_{i2}$ | $\tau_{i3}$ | $\tau_{i4}$ | $\tau_{i5}$ | $\tau_{i6}$ |
| --- | --- | --- | --- | --- | --- | --- |
| **1** | 0.0012<br>(0.0001) | 1.44<br>(0.21) | 0.47<br>(0.01) | 0.13<br>(0.02) | 0.006<br>(0.000) | 0.11<br>(0.03) |
| **2** | 0.0013<br>(0.0001) | 1.37<br>(0.21) | 0.47<br>(0.01) | 0.12<br>(0.01) | 0.006<br>(0.000) | 0.05<br>(0.02) |
| **3** | 0.0014<br>(0.0001) | 1.46<br>(0.21) | 0.48<br>(0.01) | 0.13<br>(0.02) | 0.006<br>(0.000) | 0.08<br>(0.03) |
| **4** | 0.0014<br>(0.0001) | 1.24<br>(0.19) | 0.47<br>(0.01) | 0.14<br>(0.02) | 0.006<br>(0.000) | 0.10<br>(0.03) |
| **5** | 0.0013<br>(0.0001) | 1.13<br>(0.16) | 0.47<br>(0.01) | 0.14<br>(0.02) | 0.006<br>(0.000) | 0.09<br>(0.02) |
| **6** | 0.0013<br>(0.0001) | 1.55<br>(0.23) | 0.48<br>(0.01) | 0.11<br>(0.01) | 0.006<br>(0.000) | 0.11<br>(0.03) |
| **7** | 0.0015<br>(0.0001) | 1.44<br>(1.22) | 0.49<br>(0.01) | 0.12<br>(0.01) | 0.006<br>(0.000) | 0.10<br>(0.03) |
| **8** | 0.0014<br>(0.0001) | 1.06<br>(0.15) | 0.47<br>(0.01) | 0.13<br>(0.02) | 0.006<br>(0.000) | 0.08<br>(0.02) |
| **9** | 0.0015<br>(0.0001) | 1.48<br>(0.22) | 0.48<br>(0.01) | 0.10<br>(0.01) | 0.006<br>(0.000) | 0.08<br>(0.02) |
| **10** | 0.0013<br>(0.0001) | 1.51<br>(0.22) | 0.47<br>(0.01) | 0.11<br>(0.01) | 0.006<br>(0.000) | 0.07<br>(0.02) |
| **11** | 0.0014<br>(0.0001) | 1.46<br>(0.22) | 0.47<br>(0.01) | 0.11<br>(0.01) | 0.006<br>(0.000) | 0.14<br>(0.04) |
| **12** | 0.0014<br>(0.0001) | 1.40<br>(0.22) | 0.48<br>(0.01) | 0.13<br>(0.02) | 0.006<br>(0.000) | 0.06<br>(0.02) |
| **13** | 0.0014<br>(0.0001) | 1.61<br>(0.25) | 0.47<br>(0.01) | 0.12<br>(0.01) | 0.006<br>(0.000) | 0.08<br>(0.02) |

**Table S11.** Mean and 94% HWCI of the *θ_ik_* distributions (Simulation study).

| $i$ | $\theta_{ij}(1)$ | $\theta_{ij}(2)$ | $\theta_{ij}(3)$ | $\theta_{ij}(4)$ | $\theta_{ij}(5)$ | $\theta_{ij}(6)$ |
| --- | --- | --- | --- | --- | --- | --- |
| **1** | 0.0012<br>(0.0001) | 1.48<br>(0.09) | 0.46<br>(0.01) | 0.13<br>(0.00) | 0.006<br>(0.000) | 0.14<br>(0.03) |
| **2** | 0.0013<br>(0.0001) | 1.33<br>(0.09) | 0.47<br>(0.01) | 0.12<br>(0.00) | 0.006<br>(0.000) | 0.03<br>(0.03) |
| **3** | 0.0014<br>(0.0001) | 1.53<br>(0.12) | 0.48<br>(0.01) | 0.15<br>(0.00) | 0.006<br>(0.000) | 0.08<br>(0.01) |
| **4** | 0.0014<br>(0.0001) | 1.09<br>(0.07) | 0.46<br>(0.01) | 0.17<br>(0.00) | 0.006<br>(0.000) | 0.11<br>(0.01) |
| **5** | 0.0013<br>(0.0001) | 0.88<br>(0.05) | 0.47<br>(0.01) | 0.16<br>(0.01) | 0.006<br>(0.000) | 0.08<br>(0.01) |
| **6** | 0.0013<br>(0.0001) | 1.77<br>(0.10) | 0.47<br>(0.01) | 0.11<br>(0.01) | 0.006<br>(0.000) | 0.13<br>(0.01) |
| **7** | 0.0016<br>(0.0001) | 1.49<br>(0.09) | 0.49<br>(0.01) | 0.11<br>(0.01) | 0.006<br>(0.000) | 0.10<br>(0.01) |
| **8** | 0.0014<br>(0.0001) | 0.80<br>(0.04) | 1.48<br>(0.01) | 0.14<br>(0.00) | 0.006<br>(0.000) | 0.07<br>(0.01) |
| **9** | 0.0017<br>(0.0001) | 1.57<br>(0.07) | 1.50<br>(0.01) | 0.08<br>(0.00) | 0.006<br>(0.000) | 0.07<br>(0.01) |
| **10** | 0.0012<br>(0.0001) | 1/66<br>(0.09) | 0.46<br>(0.01) | 0.10<br>(0.00) | 0.006<br>(0.000) | 0.05<br>(0.01) |
| **11** | 0.0014<br>(0.0001) | 1.51<br>(0.07) | 0.48<br>(0.01) | 0.11<br>(0.00) | 0.006<br>(0.000) | 0.22<br>(0.01) |
| **12** | 0.0014<br>(0.0001) | 1/43<br>(0.11) | 0.47<br>(0.01) | 0.14<br>(0.00) | 0.006<br>(0.000) | 0.04<br>(0.01) |
| **13** | 0.0014<br>(0.0001) | 1.82<br>(0.12) | 0.48<br>(0.01) | 0.11<br>(0.00) | 0.006<br>(0.000) | 0.07<br>(0.01) |

**Table S12.** RMSE, HWCI and *R*^2^ for the z-score learning curves at the population and test levels.

| | | RMSE | HWCI | $R^2$ |
| --- | --- | --- | --- | --- |
| Population | Congruent, low contrast | 0.014 | 0.036 | 0.994 |
|  | Incongruent, low contrast | 0.003 | 0.019 |  |
|  | Congruent, medium contrast | 0.015 | 0.038 |  |
|  | Incongruent, medium contrast | 0.006 | 0.022 |  |
|  | Congruent, high contrast | 0.014 | 0.036 |  |
|  | Incongruent, high contrast | 0.007 | 0.051 |  |
| Test | Congruent, low contrast | 0.010 | 0.033 | 0.994<br>(0.003) |
|  | Incongruent, low contrast | 0.007 | 0.017 |  |
|  | Congruent, medium contrast | 0.011 | 0.035 |  |
|  | Incongruent, medium contrast | 0.012 | 0.022 |  |
|  | Congruent, high contrast | 0.010 | 0.034 |  |
|  | Incongruent, high contrast | 0.012 | 0.052 |  |

**Table S13.** RMSE, HWCI and *R*^2^ for the *d*’ learning curves at the population and test levels.

| | | RMSE | HWCI | $R^2$ |
| --- | --- | --- | --- | --- |
| Population | Low contrast | 0.037 | 0.021 | 0.981 |
|  | Normal contrast | 0.077 | 0.050 |  |
|  | High contrast | 0.120 | 0.084 |  |
| Test | Low contrast | 0.041 | 0.020 | 0.986<br>(0.003) |
|  | Normal contrast | 0.073 | 0.046 |  |
|  | High contrast | 0.090 | 0.077 |  |

**Table S14.** RMSE, HWCI and *R*^2^ for the z-score learning curves in the prediction tasks.

| | | RMSE | HWCI | $R^2$ |
| --- | --- | --- | --- | --- |
| No data | Congruent, low contrast | 0.010 | 0.142 | 0.972 |
|  | Incongruent, low contrast | 0.037 | 0.076 |  |
|  | Congruent, medium contrast | 0.010 | 0.150 |  |
|  | Incongruent, medium contrast | 0.026 | 0.089 |  |
|  | Congruent, high contrast | 0.014 | 0.146 |  |
|  | Incongruent, high contrast | 0.017 | 0.218 |  |
| 300 trials | Congruent, low contrast | 0.022 | 0.095 | 0.968 |
|  | Incongruent, low contrast | 0.028 | 0.051 |  |
|  | Congruent, medium contrast | 0.020 | 0.100 |  |
|  | Incongruent, medium contrast | 0.028 | 0.072 |  |
|  | Congruent, high contrast | 0.012 | 0.100 |  |
|  | Incongruent, high contrast | 0.019 | 0.142 |  |
| 2700 trials | Congruent, low contrast | 0.016 | 0.060 | 0.980 |
|  | Incongruent, low contrast | 0.028 | 0.042 |  |
|  | Congruent, medium contrast | 0.013 | 0.063 |  |
|  | Incongruent, medium contrast | 0.023 | 0.042 |  |
|  | Congruent, high contrast | 0.007 | 0.064 |  |
|  | Incongruent, high contrast | 0.014 | 0.083 |  |
| 9600 trials | Congruent, low contrast | 0.014 | 0.033 | 0.994 |
|  | Incongruent, low contrast | 0.009 | 0.012 |  |
|  | Congruent, medium contrast | 0.012 | 0.034 |  |
|  | Incongruent, medium contrast | 0.010 | 0.025 |  |
|  | Congruent, high contrast | 0.006 | 0.034 |  |
|  | Incongruent, high contrast | 0.012 | 0.051 |  |

**Table S15.** RMSE, 94% HWCI and *R*^2^ for the *d*’ learning curves in the prediction tasks.

| | | RMSE | HWCI | $R^2$ |
| --- | --- | --- | --- | --- |
| No data | Low contrast | 0.112 | 0.077 | 0.960 |
|  | Medium contrast | 0.096 | 0.184 |  |
|  | High contrast | 0.105 | 0.317 |  |
| 300 trials | Low contrast | 0.132 | 0.065 | 0.942 |
|  | Medium contrast | 0.150 | 0.146 |  |
|  | High contrast | 0.115 | 0.226 |  |
| 2700 trials | Low contrast | 0.111 | 0.038 | 0.966 |
|  | Medium contrast | 0.114 | 0.083 |  |
|  | High contrast | 0.078 | 0.133 |  |
| 9600 trials | Low contrast | 0.059 | 0.025 | 0.987 |
|  | Medium contrast | 0.065 | 0.055 |  |
|  | High contrast | 0.067 | 0.081 |  |

## Footnotes

1 Although the BIP and HBM are formulated to model trial-by-trial data, we use *t* to denote each block of 300 trials in the rest of the paper.

